# Voxel-accurate MRI-microscopy correlation enables AI-powered prediction of brain disease states

**DOI:** 10.1101/2025.10.06.680637

**Authors:** Julian Schroers, Yvonne Yang, Ekin Reyhan, Nirosan Sivapalan, Einar Ismail-Zade, Alina Heuer, Jonas G. Scheck, Atefeh Pourkhalili Langeroudi, Obada T. Alhalabi, Tara Moghiseh, Manuel Fischer, Johann Jende, Bogdana Suchorska, Dieter Henrik Heiland, Matthia A. Karreman, Franz L. Ricklefs, Michael O. Breckwoldt, Felix T. Kurz, Varun Venkataramani

## Abstract

Magnetic resonance imaging (MRI) is essential for visualizing the healthy and diseased brain, yet the cellular basis of MRI signal and how it changes over time remain poorly understood. Here, we present **BRIDGE** (**B**rain **R**adiological **I**maging with **D**eep-learning based **G**round-Truth **E**xploration), a platform integrating *in vivo* MRI with *in vivo* two-photon (2P) and *ex vivo* super-resolution microscopy using a multi-step, iterative co-registration pipeline. It enables *in vivo*, longitudinal, and voxel-precise mapping of MRI signals to their cellular origins for the first time. The registered overlay reveals the cellular and anatomical origins of MRI signals and enables training of convolutional neural networks to enhance the effective resolution of MRI. Using BRIDGE, we identified a microenvironmental vessel biomarker for early metastatic colonization in patient-derived xenograft models of brain metastasis. In particular we found that distinct T2*-weighted hypointense lesions correspond to reduced blood flow and erythrostasis in perimetastatic capillaries. In glioma, longitudinal intravital studies further demonstrated direct correlations between non-vasogenic T2-weighted signal changes and patient-dependent tumor growth dynamics. Taken together, BRIDGE advances radiological interpretation by establishing a microscopic ground truth for MRI signatures over time, enabling deep learning-based predictive histology, and providing cellular-level insights into tumor microenvironment features with direct clinical imaging implications.

**Graphical abstract:** **BRIDGE enables longitudinal voxel-to-voxel correlation and ground truth based automatic segmentation of MR images**

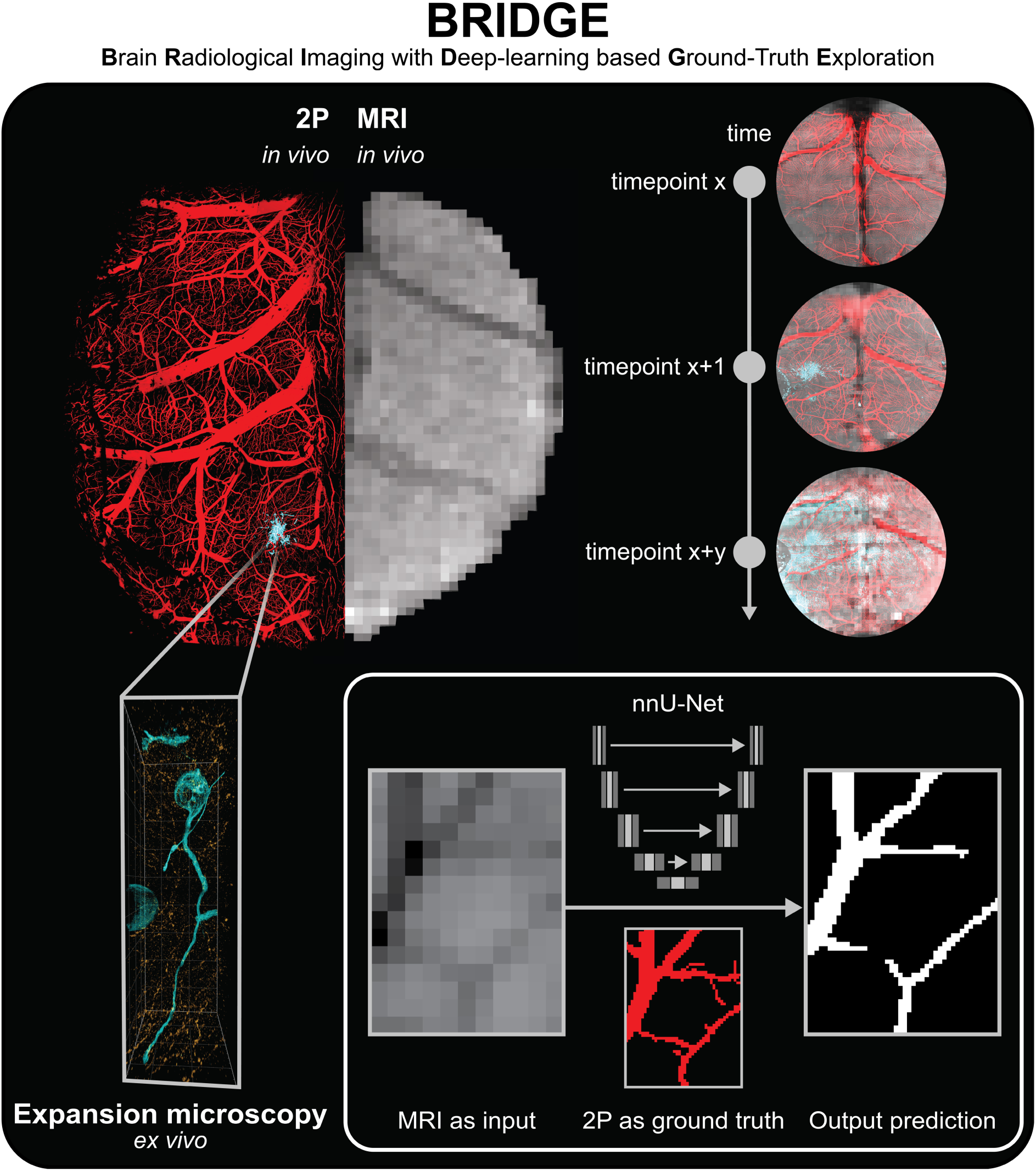

## Introduction

Magnetic resonance imaging (MRI) is an important radiological technology to visualize structural and anatomical abnormalities and disease processes [1]. Core sequences, such as T1-weighted (T1w), T2-weighted (T2w), and T2*-weighted (T2*w), are essential in diagnosing and monitoring neurological conditions, particularly those involving brain vasculature and tumors [2–4]. Radiological assessments rely on qualitative evaluations of visible abnormalities, introducing subjective variability due to the expertise of practitioners [4–7]. The emergence of quantitative radiomics, driven by multimodal data analysis, interrater segmentation evaluations, and quantitative MRI, introduced a more objective evaluation of imaging signatures [8–10]. More recently, artificial neural networks have been applied to tasks such as predictive modeling and synthetic sequence generation, offering new tools for clinical diagnosis [11, 12].

However, current AI-assisted predictions rely on manual human segmentation, which can be variable and lacks robust validation [8, 13]. Therefore, an accurate correlation of microscopic cellular ground truth with MRI signal is needed.

Recent studies have started to simultaneously acquire MRI and microscopy images [14–20]. However, existing pipelines for correlative imaging using two-photon (2P) microscopy and MRI either did not directly overlay microscopy and MRI modalities [14, 15] or were not able to provide a voxel-accurate multimodal registration [16]. While light-sheet microscopy enables the investigation of whole mouse brain volumes, it is limited to static, single-time-point measurements at microscopic resolution [17, 21–23]. Therefore, the need for longitudinal studies capable of tracking changes in single voxels over time with 3D spatial information and precise image co-registration remains unmet. Lastly, it remains unclear whether microscopy-based ground truth co-registered with MRI could enable AI-driven prediction of cellular structures directly from MRI data.

To address these limitations, we developed BRIDGE (**B**rain **R**adiological **I**maging with **D**eep-learning based **G**round-Truth **E**xploration), a pipeline enabling comprehensive voxel-to-voxel accurate co-registration between multimodal *in vivo* MRI, intravital 2P microscopy, and *ex vivo* light and super-resolution microscopy in the healthy brain as well as patient-derived brain tumor models, including glioblastoma and brain metastases. We acquired 126 correlative datasets with BRIDGE, including patient-derived xenograft models and resected tissue from human patients across various MRI sequences and 3D scales.

BRIDGE can be used in a variety of ways. First to enhance structural detection and subsequent automatic segmentation, BRIDGE utilizes microscopy data as ground truth to train MR images with convolutional neural networks [24, 25]. This approach paves the way to improve the effective resolution of MRI with microscopy.

Moreover, BRIDGE is suitable for understanding the biological processes involved in neurological diseases and their effects on MRI signal. For this purpose, we investigated brain metastases and glioblastoma. Previously in glioblastoma, attempts have been made to attribute biological significance to T2 signal: Preceding research has demonstrated that non-local tumor progression is associated with T2-hyperintense, non-contrast-enhancing tumor regions [26]. This has led to the hypothesis that T2w hyperintensity could be an early indicator of glioma progression. However, it is still controversial how exactly temporal changes in T2/FLAIR imaging are related to disease progression and if they could guide clinical decisions [27].

In particular, the differentiation of T2-positive areas in glioma between edema and tumor progression is still challenging [28, 29]. Importantly, these observations lack histopathological correlation and biological ground truth [30]. Therefore, BRIDGE enables to correlate glioblastoma dynamics with MRI signal changes and translate these findings to the human disease.

## Results

### Multimodal imaging co-registration bridges MRI and *in* and *ex vivo* light microscopy resolution

Accurate co-registration between intravital two-photon (2P) microscopy and magnetic resonance imaging (MRI) is essential for linking microscopic cellular structures with macroscopic imaging data *in vivo*. For this purpose, we established the BRIDGE pipeline **(Figure 1)** consisting of repetitive longitudinal correlative 2P and 9.4T high-resolution MRI acquisitions. Subsequently, *ex vivo* microscopy, including expansion microscopy, was performed to acquire correlated, super-resolved light microscopic data after transcardial perfusion. Next, multimodal and longitudinal registration requiring precise image adjustments such as an elastic registration method based on vessel bifurcations were iteratively performed. Voxel-to-voxel-registered data, data normalization, ROI selection, and statistical evaluation were used for downstream analyses and interpretation. Finally, 2P data provided the foundation for a deep-learning-based restoration of MR images, enhancing resolution and structural detail. **(Figure 1A)**

**Figure 1.**
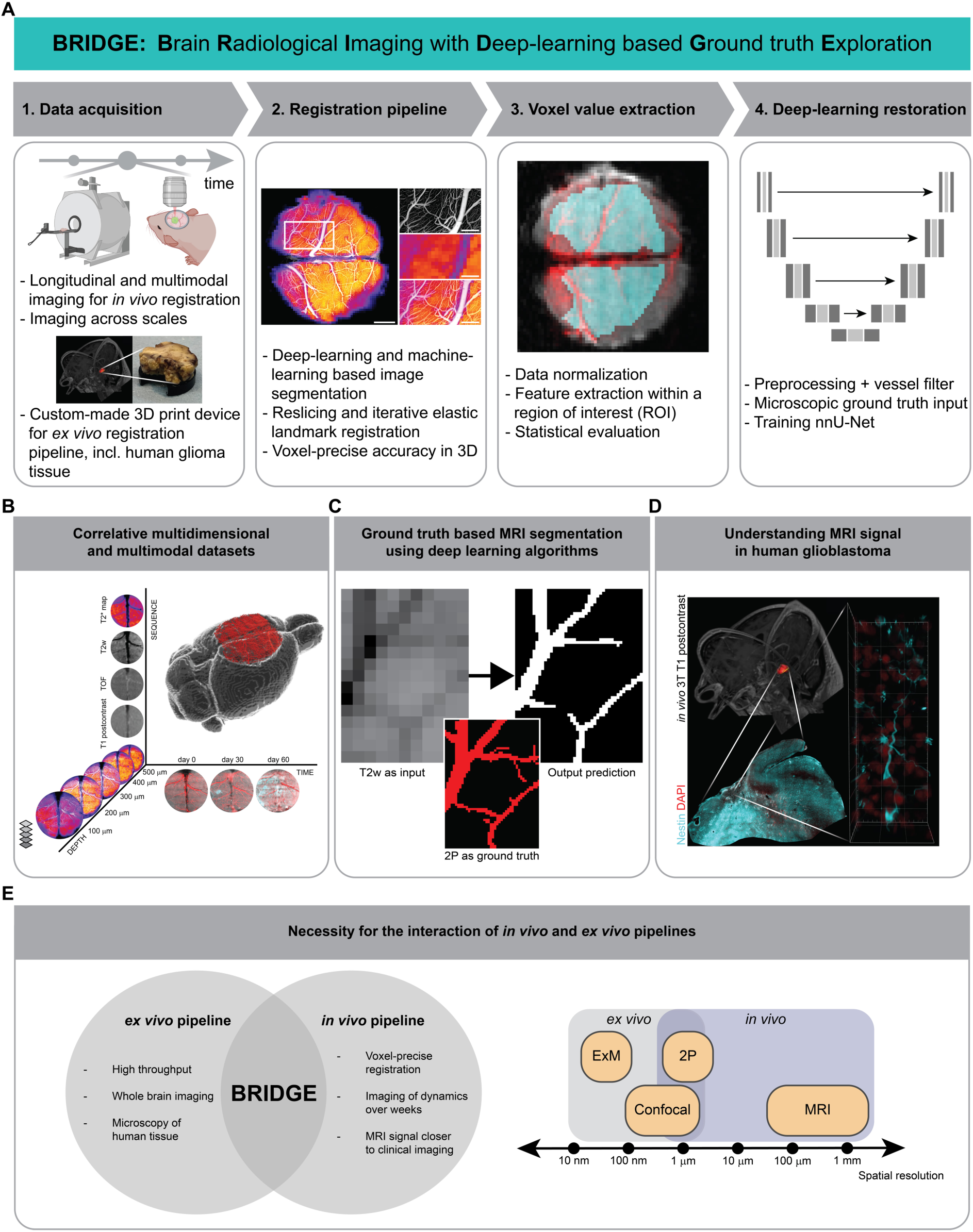
Overview of the BRIDGE pipeline A, The BRIDGE pipeline consists of 4 major steps: Data acquisition, registration pipeline, voxel value extraction and deep-learning based image restoration. **B,** Multidimensional correlation of MRI and 2P microscopy showing imaging depth (z-axis), MRI sequences (y-axis), and time (x-axis). **C,** Illustration of ground truth-based MRI segmentation using deep learning algorithms. **D,** Illustration of understanding MRI signal in human glioblastoma. **E,** Left: Integration of *in vivo* and *ex vivo* pipelines using BRIDGE. Right: Spatial resolutions of imaging techniques subdivided into *in vivo* and *ex vivo* imaging.

The multidimensional output of BRIDGE included a variety of MRI sequences (T1 native, T1 post-contrast, TOF, T2w, T2*w and T2* map), the temporal resolution provided by multiple measurements over time, and the increased imaging resolution and depth achieved through 2P which allows for improved tissue penetration **(Figure 1B, Movie S1)**.

Additionally, BRIDGE enhanced structural detection by leveraging microscopy as a reference to train MR images using a convolutional neural network for image segmentation [24] **(Figure 1C)**.

As a first step toward clinical translation, we demonstrated how *in vivo* MRI of glioblastoma patients could be directly correlated with confocal and super-resolution microscopy **(Figure 1D)**.

A crucial part of this work is the integration of our *in vivo* and *ex vivo* co-registration and analysis pipelines. While the *in vivo* pipeline could track subcellular dynamics over weeks, the *ex vivo* pipeline enables high-throughput endpoint imaging of human and mouse tissue combined with whole brain imaging and super-resolution microscopy. The interface between *in vivo* and *ex vivo* imaging combines high temporal resolution with high spatial resolution enabling to bridge scales. **(Figure 1E)**

### Longitudinal imaging across scales with voxel-accurate registration

BRIDGE combines longitudinal *in vivo* MRI and 2P microscopy through repetitive acquisitions followed by transcardial perfusion for brain tissue extraction and *ex vivo* correlative MRI studies with *ex vivo* expansion super-resolution microscopy [31–36] **(Figure 2A)**. To reduce MRI artifacts and enable an accurate registration, custom-made teflon rings instead of conventional titanium rings were used [15] **(Figures S1A-B)**.

**Figure 2.**
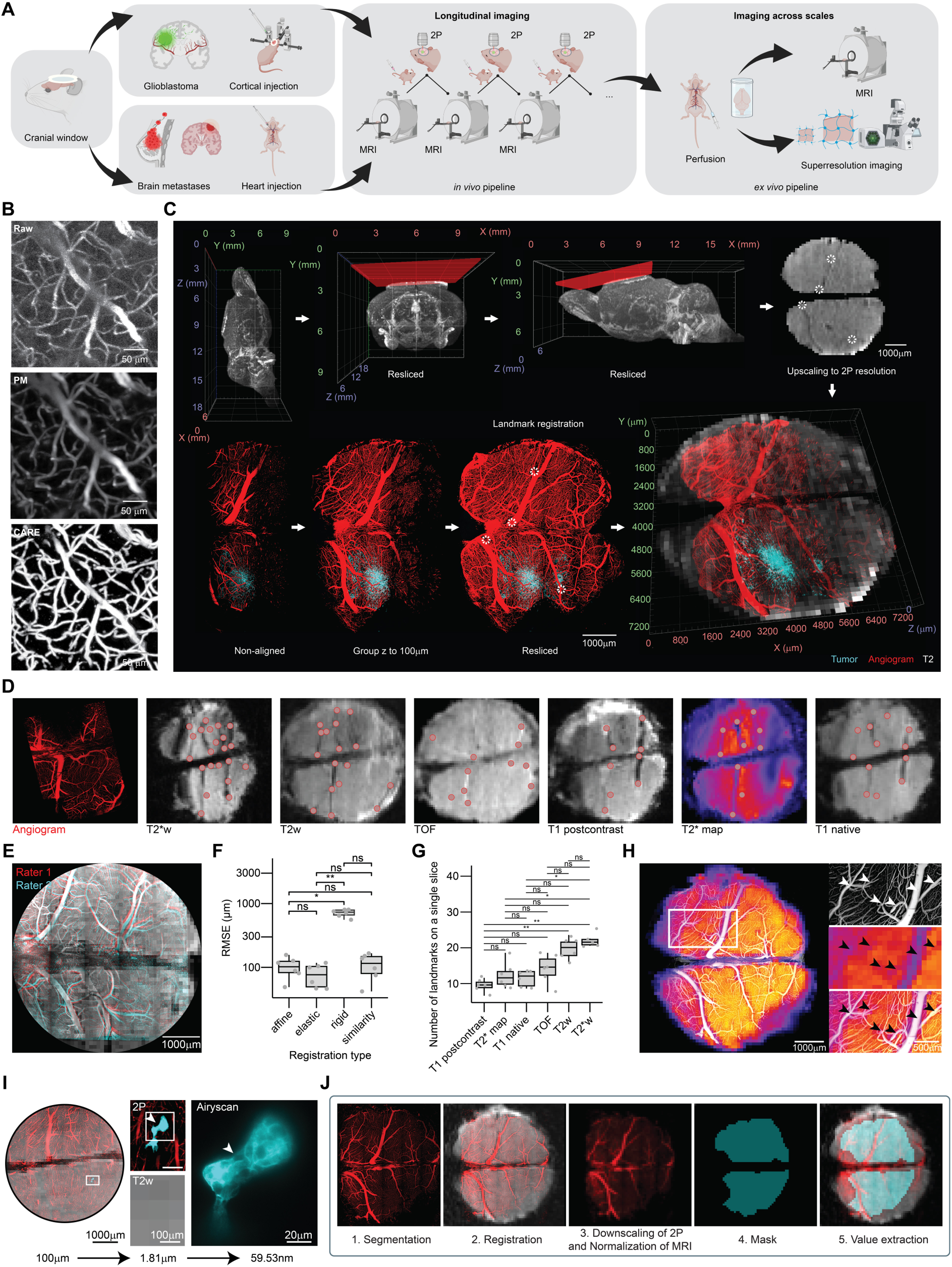
Longitudinal voxel-accurate correlative MRI and intravital 2P microscopy reveals structural and functional changes over time. **A**, Schematic of image acquisition for BRIDGE. **B**, Content-aware restoration (CARE) of 2P microscopy and subsequent interactive machine learning segmentation. **C**, Multistep and iterative registration workflow including reslicing MRI and 2P images, rescaling, and elastic 3D landmark registration using blood vessel landmarks. **D,** Comparison of vessel branching points as possible landmarks regarding different sequences in an exemplary dataset. **E**, Example of inter-rater evaluation. **F**, RMSE comparison across registration modes. Median with interquartile range (n = 6 datasets per registration mode, Kruskal-Wallis test followed by Dunn-Bonferroni post-hoc test). **G**, Number of identified landmarks across MRI sequences. Median with interquartile range (n = 6 datasets per sequence, Kruskal-Wallis test followed by Dunn-Bonferroni post-hoc test). **H**, T2* map and 2P-based angiogram demonstrating the precision of the pipeline. Arrowheads indicate registration accuracy of small vessels**. I,** *in vivo* imaging across scales of Jimt1 metastasis with Airyscan two-photon microscopy. **J,** Visualization of the voxel intensity extraction workflow of the co-registered MRI and 2P images.

Acquiring cortex-wide tile scans with *in vivo* 2P microscopy of the brain vasculature using a fluorescent angiogram is crucial for bridging macroscopic MRI resolution with the microscopic resolution as uniquely identifiable vessels and vessel branching points serve as landmarks. However, achieving broad spatial coverage requires rapid imaging as anesthesia time of mice is limited [37] which can compromise image resolution and signal-to-noise-ratio (SNR). Here, we adapted our 2P imaging workflow to incorporate deep-learning-based content-aware restoration [25] and interactive machine learning using the auto context algorithm [38, 39] to enhance both SNR and resolution across the spatial imaging areas between 6.30×10^9^ μm^3^ and 2.11×10^10^ μm^3^ **(Figure 2B)**.

For cross-modality registration, preprocessed 2P and MR images undergo signal intensity normalization and multiple, iterative registration steps in BRIDGE including reslicing and an elastic, landmark-based registration method **(Figure 2C, Figure S1C-D)**. Microscopy data could be correlated across different MRI sequences **(Figure 2D)**. The registration accuracy of BRIDGE was validated through inter-rater evaluations, achieving an average root mean square error (RMSE) of 71.6 µm **(Figure 2E)**.

We evaluated four different registration algorithms that included affine, elastic, rigid, and similarity registration. Elastic registration showed a tendency towards being the most precise registration **(Figure 2F)**. Furthermore, we assessed different MR sequences for the number of visible anatomical registration landmarks and found that both T2w and T2*w sequences [40, 41] had the highest number of identifiable registration points in 3D **(Figures 2D, G)**. When correlating 2P images to T2* maps, even small vessels with a diameter down to approximately 40 µm showed correspondence between the angiogram and T2* visible vessels **(Figure 2H, Movie S1)**.

Using BRIDGE and different modalities of 2P imaging, the workflow successfully correlated 9.4T MRI to super-resolution Airyscan imaging [42, 43] *in vivo* **(Figure 2I)**. Further, it is applicable to various biological models, such as patient-derived brain tumor models **(Figure S1E, Movie S1)**. For subsequent quantitative analyses, the resolution of the 2P images was downsampled and matched to the resolution of the MR images (see Materials and Methods). Subsequently, the corresponding voxel intensities were extracted from the registered 2P and MRI data within a region of interest (ROI) **(Figure 2J)**.

### *Ex vivo* expansion super-resolution microscopy-MRI correlation enhances effective image resolution of macroscopic imaging

To further push the limits of image resolution with correlative MRI technologies, we integrated an *ex vivo* microscopy pipeline using expansion microscopy as a super-resolution microscopy approach [31–36]. To ensure a cross-modality, standardized reference plane, we developed a custom 3D reslicing device for vibratome sectioning **(Figure 3A)**. This device is specifically designed to mount brain tissue on a pedestal serving as a common orientation plane for *ex vivo* MRI as well as subsequent slicing and microscopy **(Figure 3B)**. It includes a lunar notch designed to prevent air bubble accumulation near the brain tissue, thereby reducing the risk of introducing artifacts into the MR images [44] **(Figure S2B)**. Furthermore, the stable mounting brings the *ex vivo* tissue closer to the MRI coil, enhancing the SNR [45].

**Figure 3.**
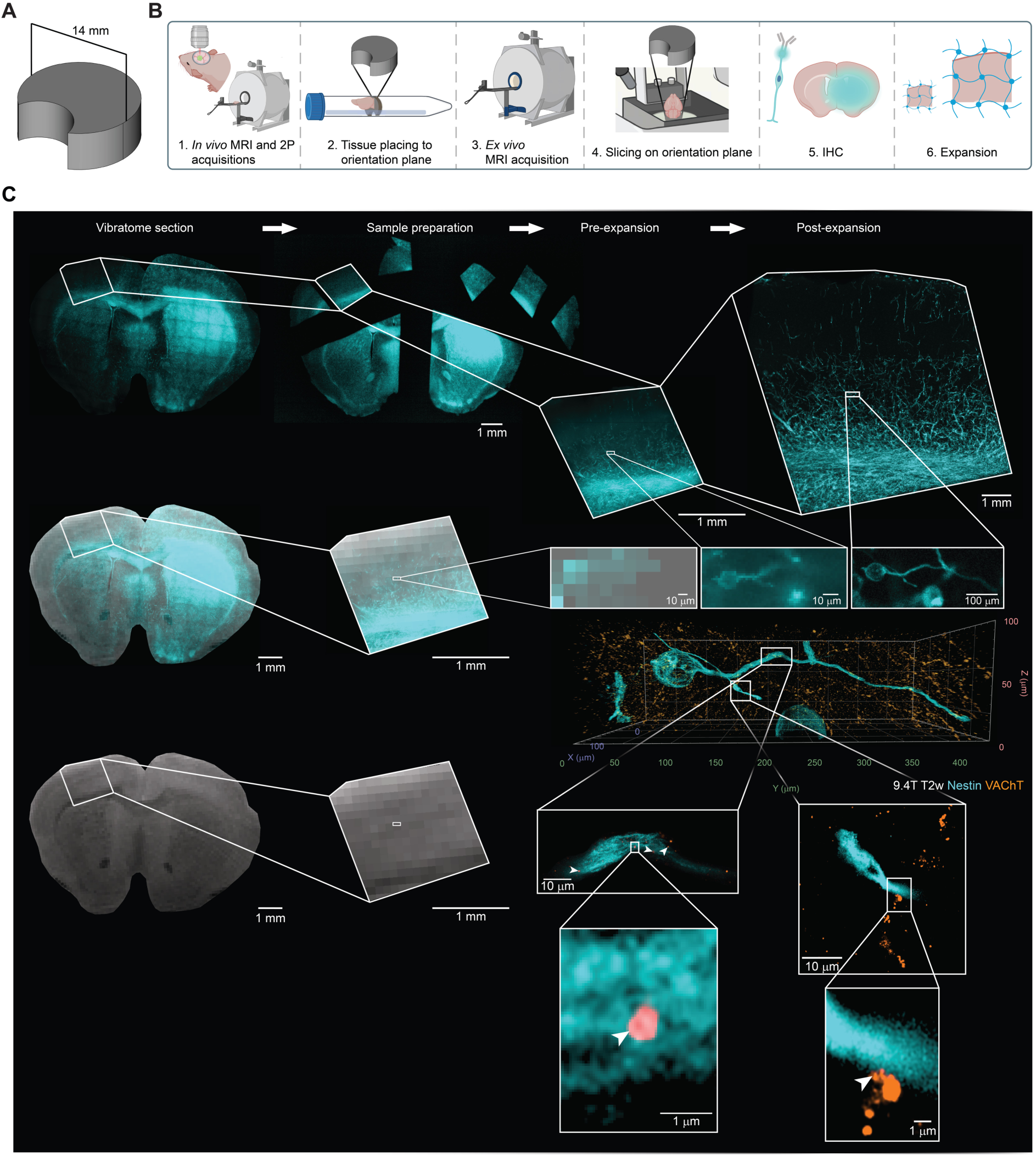
Correlative MRI and super-resolution microscopy *ex vivo*. **A**, 3D model of the custom device used in the *ex vivo* correlation pipeline. **B**, Schematic of the patient-derived mouse model image acquisition pipeline across scales. **C**, Correlative *ex vivo* expansion microscopy and MRI in the patient-derived S24 glioblastoma model mouse brain. Nestin (cyan) is used as a tumor marker of glioblastoma cells and vesicular acetylcholine transporter (VAChT) is a vesicular marker of cholinergic synaptic vesicles (orange). Arrowheads depict putative neuron-to-glioma synapses (NGS).

Subsequent co-registration utilizes anatomical landmarks, ensuring precise alignment between MRI and microscopy. Since each brain section corresponds to the thickness of an MR voxel, aligning one MR slice with one brain slice ensures that all subsequent slices are aligned **(Figure S2A)**. After co-registration, it is possible to register the brain regions onto the Allen Mouse Brain Common Coordinate Framework (CCFv3) allowing a whole-brain investigation [46] **(Figure S2C)**.

To further push the resolution boundaries on the microscopic scale to the level of single synaptic boutons, we conducted correlative expansion microscopy with an expansion factor of 4 [31]. This approach enabled the visualization of putative, individual neuron-to-glioma synapses (NGS) which were previously described as hallmark of multicellular tumor networks [47–50] **(Figure 3C, Movie S2)**. Together, this workflow enables super-resolved correlative imaging of MRI. It paves the way for the development of radiological predictors of malignant connectivity by enabling the quantification of neuron-glioma synapses [47–50] in MR voxels.

### Ground-truth-based deep learning predictions improve the detection of the brain vasculature in MRI

In the next step, we leveraged our voxel-accurate registration to investigate whether AI-based image restoration with 2P microscopy as ground truth is able to enhance the effective resolution and accuracy of automatic MR segmentation. The aim was to enhance the visibility of vessel architecture at near-microscopic resolution in MRI and to demonstrate the effectiveness of this approach.

Utilizing correlated *in vivo* 2P microscopic information as ground truth, we used the nnU-Net deep-learning architecture [24] to segment blood vessels in T2w images **(Figure 4A)**. This AI-assisted prediction enhanced the visibility of smaller vessels **(Figure 4B)**. Many vessels that were barely visible in the T2w sequence became clearly visible for human evaluators while a minority of vessels were no longer visible after training **(Figure 4C)**. In each diameter category of ground truth vessels, the estimation of the diameter by the neural network prediction is significantly better than the binarized T2w sequence **(Figure 4D)**.

**Figure 4.**
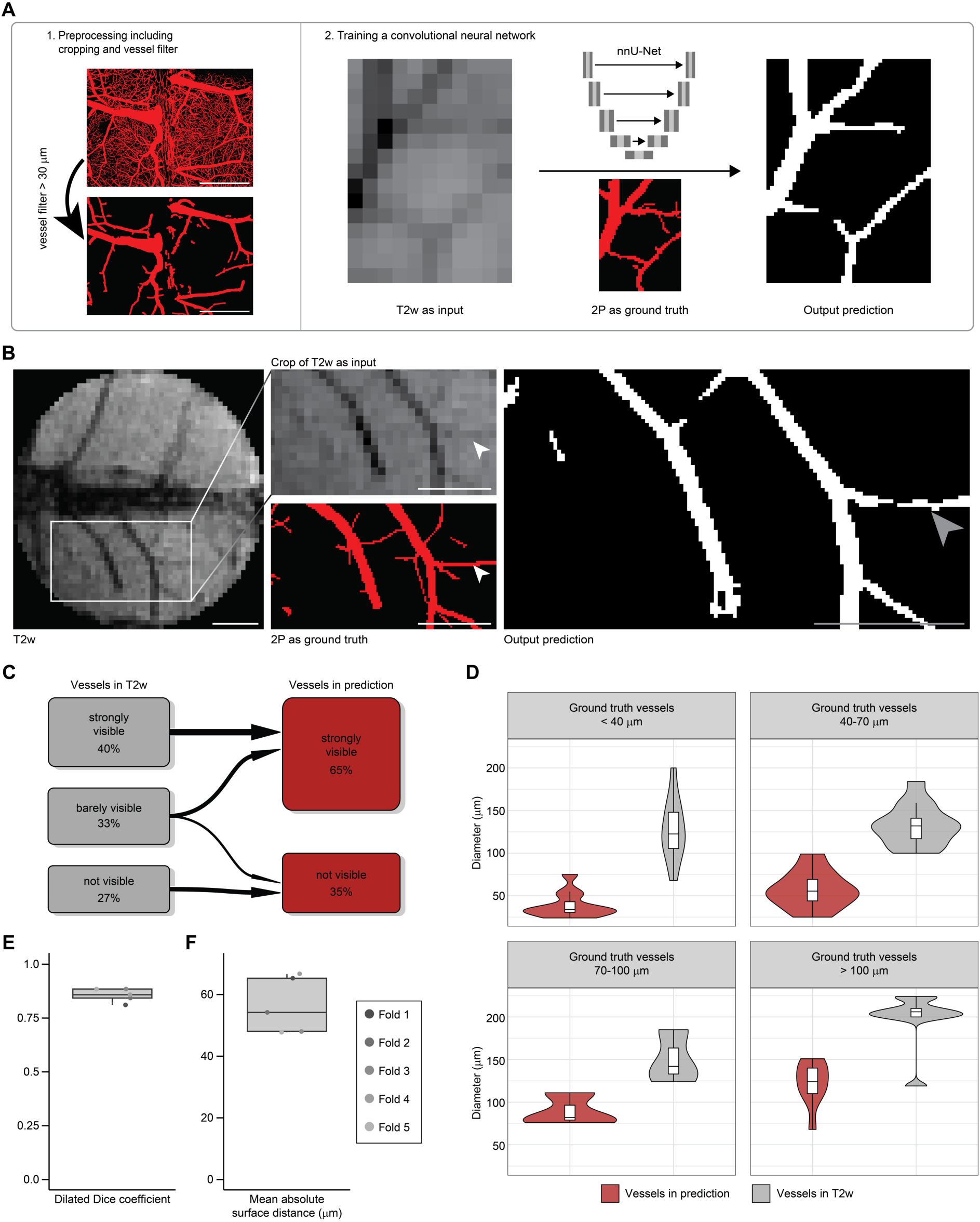
BRIDGE workflow enables ground-truth based deep learning of MRI A, Schematic of the BRIDGE training pipeline including data preprocessing and training of nnU-Net. Data preprocessing contains custom-designed vessel diameter filter. Scale bars: 1 mm. **B,** BRIDGE pipeline uses T2w images as input and 2P microscopy images as ground truth. The output is a prediction of vessels depending on the MR image and the previous training of other datasets. Arrowheads indicate small vessel which was only barely visible in T2w before training. Scale bars: 1 mm. **C,** River plot showing the vessel visibility in the T2w images and in the predictions by the neural network (n = 6 validation datasets, 51 vessels). **D,** Violin plots comparing the vessel diameters in predictions by the neural network and in T2w sequences. Each violin plot shows a different diameter category of vessels in 2P microscopy as ground truth (n = 6 validation datasets, 51 vessels). **E,** Boxplot shows Dilated Dice coefficient between ground truth and prediction. Median with interquartile range (n = 5 folds). **F,** Boxplot shows mean absolute surface distance (MASD) between ground truth and prediction. Median with interquartile range (n = 5 folds).

To assess the training, we performed a 5-fold cross-validation **(Figure S3A)**. The Dilated Dice coefficient [51] is approximately 0.86 **(Figure 4E)**. The mean absolute surface distance (MASD) [52] is about 54 μm **(Figure 4F)**, which is smaller than the isotropic MRI voxel size of 100 μm. We also performed a training in which the segmentations were randomly flipped horizontally or vertically, in contrast to the MRI images, to validate the above procedure. As expected, the results of this training were poor and not able to predict vessel structures of the ground truth **(Figure S3B)**. To summarize, this approach is able to reduce human impact on segmentations and improve the visibility of blood vessels and the estimation of the vessel diameter, thereby highlighting the potential of microscopy-trained MRI.

### Real-time correlative MRI and intravital microscopy decode signatures of blood vessels in physiology and brain metastases

We applied the BRIDGE framework to analyze the brain vessels and their visibility across different MRI sequences (T2w, T2*w, TOF) [40, 41, 53–55]. Voxel information from correlated 2P and MRI data was extracted for quantitative analyses **(Figure 2J)**. Utilizing a cryogenic coil, higher spatial resolution of 40 µm in 3D T2*w sequences enabled *in vivo* visualization of microvascular details, allowing confident correlation of vessels observed in 2P imaging, down to approximately 20 µm **(Figure 5A, Figure S4A)**.

**Figure 5.**
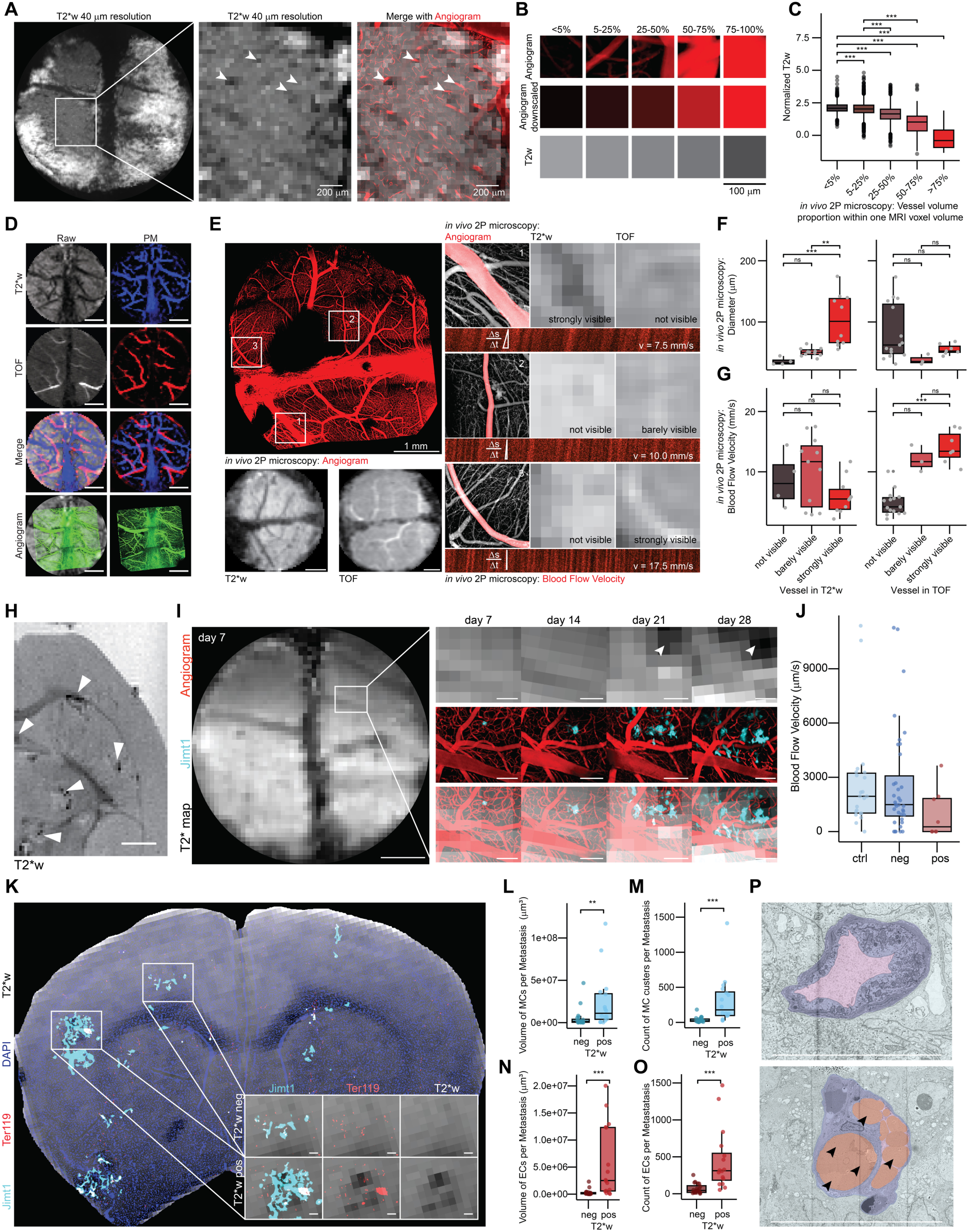
BRIDGE decodes perimetastatic T2*w changes during brain metastastic progression A, Correlation of *in vivo* high-resolution MRI (T2*w, isotropic 40 µm) with corresponding 2P microscopy angiogram. Arrowheads indicate registration accuracy of small vessels. Scale bars: 200 μm. **B,** Exemplary *in vivo* 2P-correlated MR voxels grouped based on vessel volume proportion. Scale bar: 100 μm. **C,** Vessel volume proportion plotted against normalized T2w signal intensity *in vivo*, presented as median with interquartile range (0-5%, n = 1906 voxels; 5-25%, n = 3334 voxels; 25-50%, n = 801 voxels; 50-75%, n = 196 voxels; >75%, n = 39 voxels; ANOVA with post-hoc Tukey HSD test). **D,** Probability maps of correlated *in vivo* multisequence 2P-MRI, with T2*w (blue), TOF (red), and 2P as ground truth (green). Scale bars: 1 mm. **E,** 2P intravital brain angiogram imaging mapped onto *in vivo* multisequence MRI, with examples of blood flow velocity measurements using line scans in 2P microscopy. Scale bars: 1 mm. **F,** Diameter of vessels depending on visibility across *in vivo* MRI modalities (T2*w, TOF), presented as median with interquartile range (n = 2 mice; for T2*w: not visible, n = 4 vessels; barely visible, n = 11 vessels; visible, n = 4 vessels; strongly visible, n = 6 vessels; for TOF: not visible, n = 15 vessels; barely visible, n = 3 vessels; visible, n = 4 vessels; strongly visible, n = 3 vessels; Kruskal-Wallis test followed by Dunn-Bonferroni post-hoc test). **G,** Blood flow velocity in vessels depending on visibility across *in vivo* MRI modalities (T2*w, TOF). Median with interquartile range (n values same as e, Kruskal-Wallis test followed by Dunn-Bonferroni post-hoc test). **H,** *Ex vivo* MRI screening of breast cancer brain metastasis model Jimt1 using T2*w imaging. Arrowheads show hypointense changes. Scale bar: 1 mm. **I,** Weekly *in vivo* 2P-MRI correlative imaging of an emerging T2*w hypointense Jimt1 metastasis (cyan) with angiogram (red) over 28 days. Scale bars: 1 mm (T2* map overview) and 200 µm. **J,** Blood flow velocity measurements of perimetastatic capillaries (Jimt1 mGFP in cyan) and controls with a diameter <12 μm across different *in vivo* T2* map images. Median with interquartile range (control, n = 22; T2*w negative, n = 36; T2*w positive, n = 8). **K,** *Ex vivo* MRI-LM correlation in a Jimt1 mGFP (cyan) xenografted mouse brain slice with DAPI (blue) and Ter119 (red) staining, registered onto *ex vivo* T2*w. Insets show T2*-negative and T2*-positive lesions. Scale bars: 100 µm. **L,** Comparison of total volume of metastatic cell clusters (MCs) per metastasis between T2*-negative and T2*-positive metastases *ex vivo*. Median with interquartile range (n = 3 mice; T2*-negative, n = 20; T2*-positive, n = 16; Mann-Whitney U test). **M,** Comparison of the count of MCs per metastasis between T2*-negative and T2*-positive metastases *ex vivo*. Median with interquartile range (n = 3 mice; T2*-negative, n = 20; T2*-positive, n = 16; Mann-Whitney U test). **N,** Comparison of total volume of erythrocyte clusters (EC) per metastasis between T2*-negative and T2*-positive lesions *ex vivo*. Median with interquartile range (n = 3 mice; T2*-negative, n = 20; T2*-positive, n = 16; Mann-Whitney U test). **O,** Comparison of the count of ECs per metastasis between T2*-negative and T2*-positive lesions *ex vivo*. Median with interquartile range (n = 3 mice; T2*-negative, n = 20; T2*-positive, n = 16; Mann-Whitney U test). **P,** Electron microscopy (EM) of perimetastatic capillaries showing a vessel with an empty lumen (top) and a vessel with erythrostasis and a capillary loop formation (bottom). Arrowheads indicate erythrocytes. Jimt1 (cyan) and perimetastatic capillaries with endothelium (purple), erythrocytes (red) and lumen (pink). Scale bars: 10 µm.

Voxel data were categorized based on 2P microscopy-derived blood vessel volume per voxel **(Figure 5B)**. Previous studies have noted the role of venous blood vessels in contributing to hypointense voxels in T2w and T2*w imaging [56]. Consistent with previous findings, our BRIDGE workflow demonstrated a significant inverse correlation between 2P vessel volume density per MRI voxel and T2w intensity **(Figure 5C)**. Subsequently, we assessed the visibility of blood vessels across different MRI sequences. MR-visible arteries and veins were segmented using T2* maps and Time-of-Flight (TOF) sequences [57] **(Figure 5D)**, correlating their MRI characteristics with parameters measured in 2P, such as diameter and blood flow velocity **(Figure 5E, Figure S4B)**. Although TOF signals strongly correlated with blood flow velocity, they did not correlate with vessel diameter. In contrast, T2w and T2*w sequences were effective in delineating vascular anatomy, correlating well with vessel diameter but not flow velocity. Even smaller vessels than the voxel size were visible in the MR images **(Figures 5A, F-G, Figure S4C)**. The findings support existing knowledge on TOF sequences for visualizing moving blood [58] and additionally provide further evidence through direct comparison with the microscopic ground truth.

We adopted this approach to living mice with breast cancer brain metastases. *In vivo* and *ex vivo* MRI screening of the breast cancer brain metastases xenograft model Jimt1 showed T2*w hypointense lesions **(Figure 5H, Figure S4D)**. These hypointense signature was predominantly located at the edge of the metastases, which we were able to show using high-resolution *ex vivo* MRI with a cryogenic coil [59] resulting in an isotropic voxel size of 30 µm **(Figures S4E-F)**. The number of T2*w positive lesions consistently increased over time, marking a progressive worsening in the brain metastastic landscape **(Figure 5I, Figure S4G)**. Previous findings suggested that T2*w hypointensity in brain metastases was caused by unspecific microbleedings [60].

Surprisingly, our *in vivo* BRIDGE pipeline revealed an association of T2* map-positive with a trend toward reduced blood flow velocity in perimetastatic capillaries with diameters under 12 µm **(Figure 5J)**, which corresponds to approximately twice the size of a murine erythrocyte [61]. Further, we could see a heterogeneity of this radiological signature and identified both T2* map-positive and negative metastases **(Figure 2I and Figures S4H-I)**. Utilizing the *ex vivo* BRIDGE pipeline with additional immunofluorescence stainings for metastatic cell markers and the erythrocyte marker Ter119 [62], we observed increased numbers and sizes of metastatic cell clusters (MCs) and erythrocyte clusters (ECs) in T2*w positive lesions **(Figures 5K-O)**. Quasi-correlative electron microscopy of brain metastases samples in this patient-derived xenograft model further confirmed the presence of erythrostasis within these capillaries **(Figure 5P, Figure S4J)**. Together, these findings reveal a biological mechanism distinct to microbleeds that underlies the T2*w hypointensities and allows to develop novel microscopy-inspired biomarkers for radiological detection of brain metastases.

### Voxel-to-voxel correlation of MRI and intravital microscopy reveals underlying ground truth of emerging MRI signatures of glioblastoma

To investigate the dynamic changes in MR voxel properties during glioblastoma progression with the longitudinal BRIDGE workflow, assessments were conducted with two patient-derived glioblastoma models: GG16 [63], characterized by its bulky growth **(Figures 6A-B, Figure S5B)** and S24 [47, 50, 64, 65], showing a diffuse, infiltrative growth pattern **(Figures 6C-D)**. At 60 days post-injection, GG16 tumors showed visible T2w hyperintensity, whereas no tumor was detected using high-resolution MRI in the S24 model **(Figures 6A, C)**, highlighting the heterogeneous growth patterns of these models **(Figure S5D)**. Furthermore, the overall vascular architecture does not change in the early stages of tumor colonization in the patient-derived S24 infiltrative growth model, while changes can be observed during the bulky growth of the patient-derived GG16 model **(Figures 6E-F)**.

**Figure 6.**
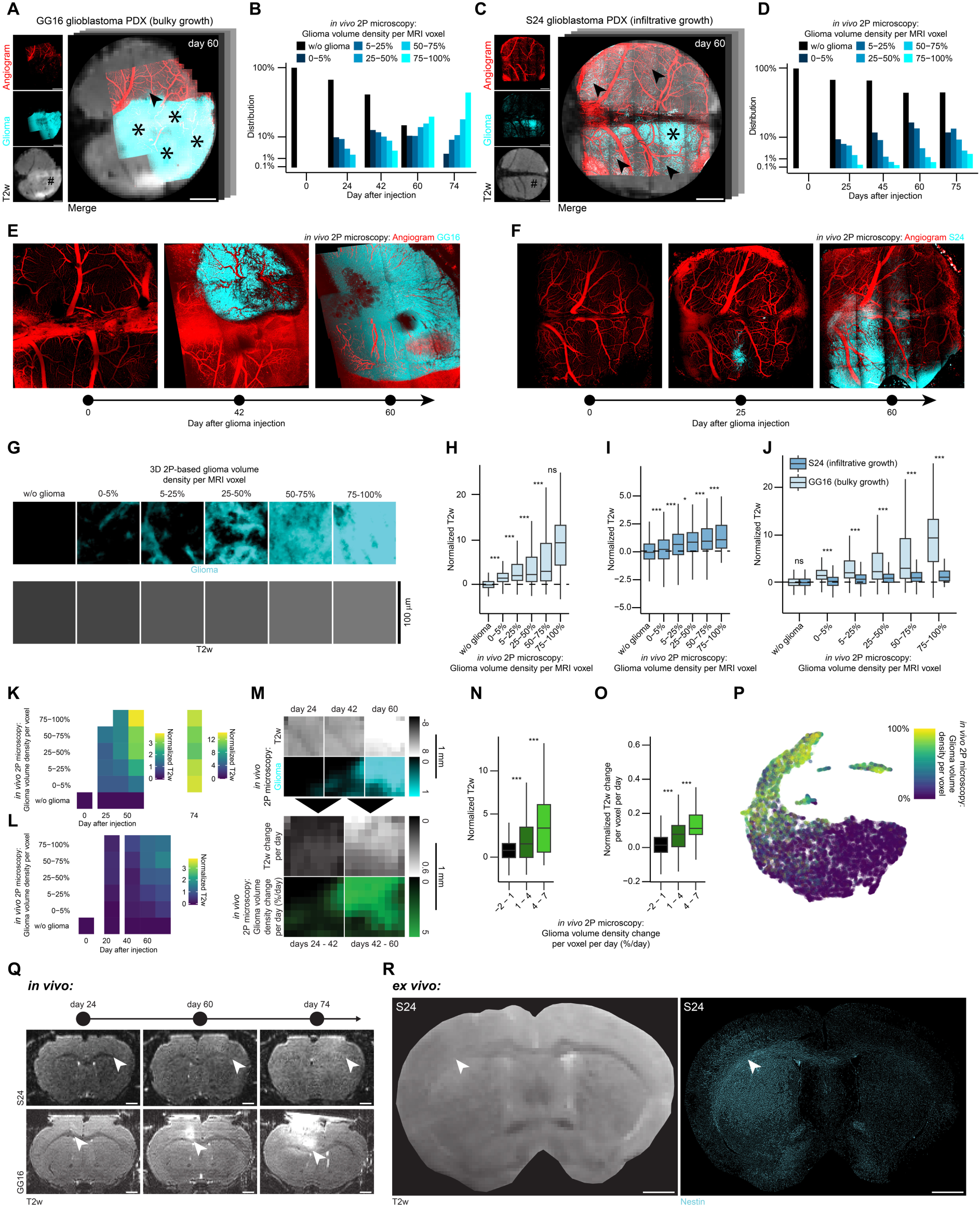
BRIDGE reveals cellular tumor burden in radiologic brain tumor features during glioblastoma progression A, *In vivo* 2P-MRI correlation of exemplary GG16 at day 60 after tumor implantation. Stars depict bulk tumor regions, arrowheads depict infiltrative regions, hashtag depicts visible tumor in T2w sequence. Scale bars: 1 mm. **B,** Barplot showing glioma volume density distribution in GG16 over time measured in 2P microscopy (n = 4 mice; day 0, n = 1 dataset, 5408 voxels; day 24, n = 3 datasets, 4636 voxels; day 42, n = 3 datasets, 7691 voxels; day 60, n = 3 datasets, 4930 voxels; day 74, n = 3 datasets, 4890 voxels). **C,** *In vivo* 2P-MRI correlation of exemplary S24 at day 60 after tumor implantation. Stars depict bulk tumor regions, arrowheads depict infiltrative regions, hashtag depicts invisible tumor in T2w sequence. Scale bars: 1 mm. **D,** Barplot showing glioma volume density distribution in S24 over time in 2P microscopy (n = 17 mice; day 0, n = 3 datasets, 11752 voxels; day 25, n = 8 datasets, 20541 voxels; day 45, n = 8 datasets, 17593 voxels; day 60, n = 11 datasets, 27689 voxels; day 75, n = 7 datasets, 18025 voxels). **E,** Timeline shows exemplary vessel architecture (red) and GG16 (cyan) at days 0, 42 and 60 after tumor implantation in the same mouse. **F,** Timeline shows exemplary vessel architecture (red) and S24 (cyan) at days 0, 25 and 60 after tumor implantation in the same mouse. **G,** Exemplary 2P-correlated MR voxels grouped based on glioma volume density with corresponding T2 voxels to visualize the correlation between glioma density and T2 signal intensity. Scale bars: 100 μm. **H,** Boxplots showing significant correlation of glioma density with normalized T2w in GG16 *in vivo*. Median with interquartile range (n = 4 mice, 13 datasets; w/o glioma, n = 14173 voxels; 0-5%, n = 2224 voxels; 5-25%, n = 2313 voxels; 25-50%, n = 2192 voxels; 50-75%, n = 2566 voxels; 75-100%, n = 4087 voxels; ANOVA with post-hoc Tukey HSD test). **I,** Boxplots showing glioma density and T2w correlation in S24 *in vivo*. Median with interquartile range (n = 17 mice, 37 datasets; w/o glioma, n = 69490 voxels; 0-5%, n = 12333 voxels; 5-25%, n = 8881 voxels; 25-50%, n = 3292 voxels; 50-75%, n = 1156 voxels; 75-100%, n = 447 voxels; ANOVA with post-hoc Tukey HSD test). **J,** Boxplots showing T2 signal intensity differences of bulky growth model GG16 and infiltrative growth model S24 *in vivo* depending on 2P microscopy glioma density per MRI voxel. Median with interquartile range (n values are the same as in f and g; Welch’s t-tests). **K,** Heatmap of glioma density compared to normalized T2w in GG16 *in vivo* (n = 4 mice, 13 datasets, 27555 voxels). **L,** Heatmap of glioma density compared to normalized T2w in S24 *in vivo* (n = 17 mice, 37 datasets, 95599 voxels). **M,** Image crops of longitudinal GG16 tumor in 2P and T2w correlation *in vivo*, highlighting voxel density changes over time. Scale bars: 1 mm. **N,** Correlation of normalized T2w signal intensity with glioma growth rate in GG16 *in vivo*. Median with interquartile range (n = 3 mice, 9 datasets; -2–1%, n = 2414 voxels; 1–4%, n = 688 voxels; 4–7%, n = 197 voxels; Kruskal-Wallis test followed by Dunn-Bonferroni post-hoc test). **O,** Change of T2w signal intensity with glioma growth rate in GG16 *in vivo*. Median with interquartile range (n values same as l, Kruskal-Wallis test followed by Dunn-Bonferroni post-hoc test). **P,** UMAP dimension reduction of normalized *in vivo* T2w, T1-native, and T1-postcontrast data for GG16, labeled by glioma volume density (n = 4 mice, 13 datasets, 27555 voxels). **Q,** Development of *in vivo* T2w signal 24, 60 and 74 days after tumor injection. Arrowheads indicate fading of CC. Top: S24, bottom: GG16. Scale bars: 1 mm. **R,** *Ex vivo* co-registration of T2w image (left) and Nestin (cyan, right) in mouse brain with S24. Arrowheads indicate fading of CC (left) and high glioma density (right). Scale bars: 1 mm.

Using our voxel value extraction workflow **(Figure 2J, Figure S5A)**, we analyzed tumor and adjacent brain parenchyma voxels **(Figure 6G)**. Both models demonstrated a significant increase in normalized T2w signal intensity corresponding to glioma volume density per MR voxel **(Figures 6H-I, Figure S5C)**, even in areas of low glioma density, typically associated with infiltrative growth [50], and in regions where manual segmentation did not detect the tumor **(Figure 6C)**. In comparison of both tumor models, the bulky growth model exhibited significantly higher normalized T2w signal intensity compared to the infiltrative model at similar glioma cell densities **(Figure 6J)**. Longitudinal analysis further showed a positive correlation of T2w signal increase with increased glioma density and time **(Figures 6K-L)**. Even though no to little vascular architecture changes occur in the infiltrative growth model S24 **(Figure 6F)**, the T2w signal intensity increases with time **(Figure 6L)**. This demonstrates that T2w signal intensity increase did not only depend on vascular architectural changes and associated vasogenic edema [66, 67], but also decodes infiltrative growth in the absence of vascular changes. Furthermore, longitudinal registration **(Figure 6M)** of GG16 datasets confirmed that normalized T2w signal intensity significantly correlated with glioma growth rate over time **(Figure 6N)**. Further, in GG16 increases in T2w signal intensity were significantly linked to glioma growth rate per time **(Figure 6O)**. To summarize, there appears to be a gradient between a T2w hyperintense infiltration zone (IZ T2+) and a T2w isointense infiltration zone (IZ T2-). This gradient seems to correlate with glioma density and glioma dynamics, i.e. growth rate and time after glioma injection.

To illustrate the correlation between MRI and 2P microscopy in the form of a smooth transition in glioma density, a UMAP dimensionality reduction [68] was employed. Information per voxel such as normalized T2w, pre- and post-contrast T1 signal intensities, and glioma volume density, effectively showing how glioma density gradations align with spatial distributions derived from multimodal MRI data **(Figure 6P, Figures S5E-J)**.

In addition to gray matter changes, we also investigated the corpus callosum during glioblastoma progression. *In vivo* imaging revealed a progressive T2w signal intensity increase of the corpus callosum over time in both S24 and GG16 **(Figure 6Q)**. Corresponding *ex vivo* microscopy and *ex vivo* T2w image confirmed that this T2 hyperintensity correlated with increased glioma cell density. Here, the infiltration zone can also be identified by a gradient in T2w imaging, ranging from IZ T2-to IZ T2+, which correlates with glioma cell density. **(Figure 6R)**

### Clinical translation of human brain tumor MRI signals using BRIDGE

Lastly, we focused on a clinical translation pipeline to apply BRIDGE to human brain tissue. For this purpose, we used a combination of *in vivo* 3T MRI, *ex vivo* 9.4T MRI and super-resolution microscopy to investigate human glioblastoma tissue which was extracted via neuronavigation **(Figures 7A-C)**. We were able to register microscopy precisely to the T2w image using vessel landmarks **(arrowheads in Figure 7B)**. These investigations confirmed that higher glioma cell density correlated with higher T2w signal intensity **(Figure 7D)**. Leveraging super-resolution microscopy, we were further able to co-register glioblastoma cells with subcellular resolution to MRI voxels. This investigation even enabled to visualize previously described neurite-like structures of glioblastoma cells that are important for the invasion into brain tissue [49, 50, 64] **(Figure 7C, Movie S3)**.

**Figure 7.**
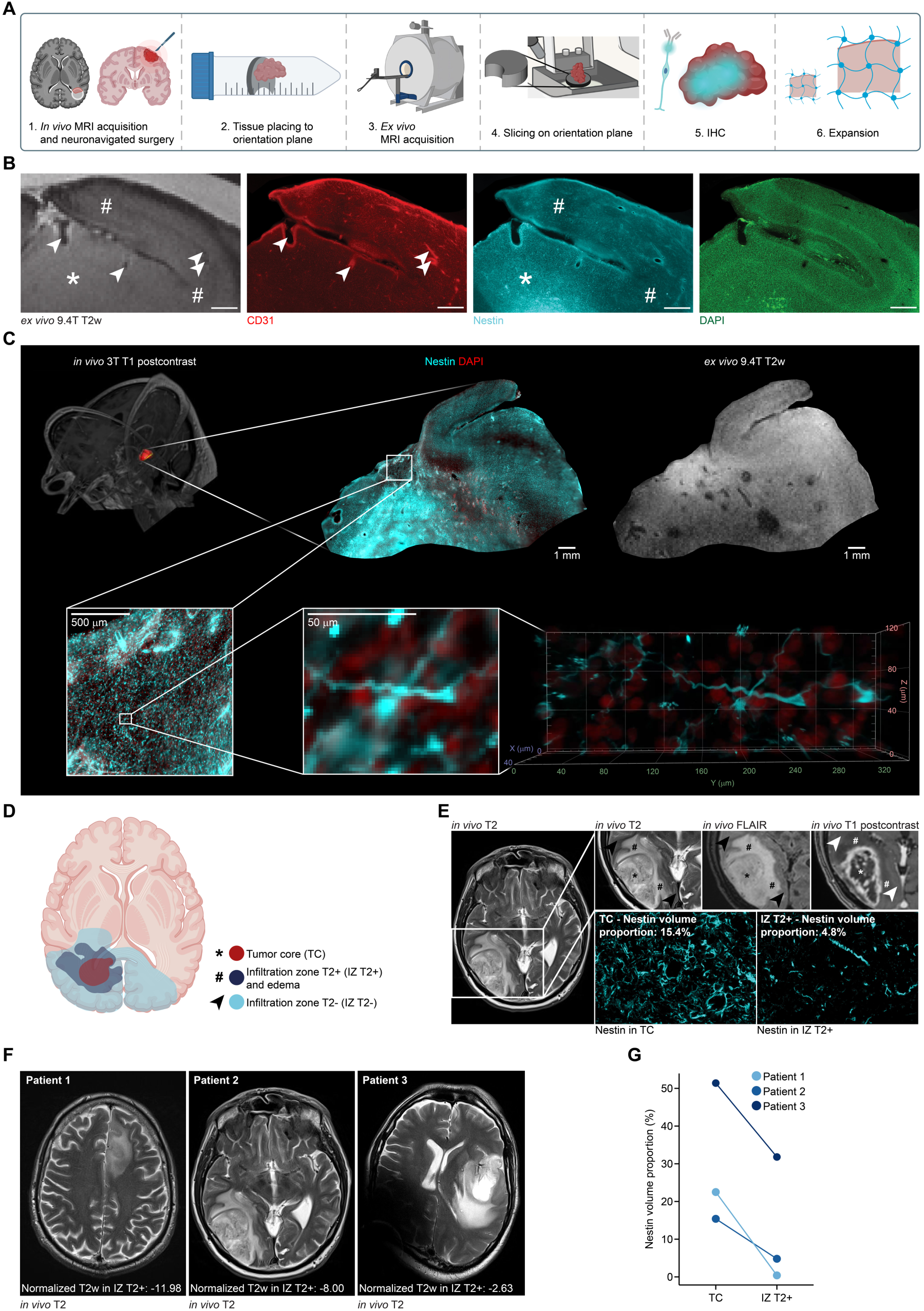
Clinical-translational investigation of human brain tumor tissue with BRIDGE A, Schematic of the human acquisition pipeline to bridge scales. **B,** *ex vivo* 9.4T T2w image as well as CD31 (red), Nestin (cyan) and DAPI (green) acquisitions. Arrowheads indicate vessels, star indicates hyperintense area with high Nestin signal and hashtags indicate isointense areas with low Nestin signal. Scale bars: 1 mm. **C,** Correlative *in vivo* and *ex vivo* MRI and *ex vivo* super-resolution microscopy in human glioblastoma tissue. Nestin (cyan) and DAPI (red). **D,** Schematic of the structure of glioblastoma, divided into TC (red), edema and IZ T2+ (dark blue) and IZ T2-(light blue). **E,** Left: Clinical T2w image of a patient with glioblastoma. Upper row shows clinical T2, FLAIR and T1 postcontrast images. TC indicates tumor core and IZ indicates infiltration zone. Stars indicate TC, hashtags indicate edema and IZ T2+ and arrowheads indicate IZ T2-. Lower row shows Nestin signal in TC with a Nestin volume proportion of 11.0% and in IZ with a Nestin volume proportion of 4.9% in the same patient. **F,** Clinical T2w images of three patients with mean normalized T2w intensity in IZ T2+. **G,** Comparison of Nestin volume proportions of the same three patients as in (F), subdivided into TC and IZ T2+, highlighting the heterogeneity of glioblastoma.

In clinical MRI, we showed that glioma densities in tumor core (TC) and IZ T2+ were highly heterogeneous in three different patients **(Figures 7D-E)**, suggesting that T2 signal also depends on glioma cell density and glioma dynamics in human patients as previously shown in our patient-derived xenograft models **(Figures 7F-G)**. Consistent with our patient-derived xenograft model data, the glioma cell density decreased from the tumor core to the infiltration zone of patients. However, the glioma cell density in the infiltration zone as well as the tumor core was patient dependent.

## Discussion

We established a framework called BRIDGE for longitudinal and multimodal voxel-accurate correlation between MRI and light microscopy, integrating *in vivo* 2P microscopy and *ex vivo* super-resolution microscopy.

Previous correlative imaging techniques have often encountered significant registration challenges, such as translation, rotation, elastic differences, and resolution adjustments. Traditional methods relying on brain slice contours as landmarks [69] or positioning the brain within pathology slice blocks [70] have shown inaccuracies, with a median distance of 390 μm between landmarks. More advanced techniques, like 3D-3D registration with CLARITY [71] still lack longitudinal registration and therefore the ability to track changes. *In vivo* correlative imaging has attempted to use vascular landmarks to align MR angiograms with microscopic imaging [18], offering higher temporal resolution, but often sacrificing spatial precision, especially in terms of axial resolution. BRIDGE addresses these limitations by improving spatial registration accuracy through an iterative registration pipeline. Our workflow provides consistent and reliable landmarks for multimodal registration using uniquely identifiable vessel branches visible in 3D on both angiograms and MRI. We evaluated MR sequences concerning the visibility of possible landmarks and found T2w and T2*w sequences to be the most suitable sequences.

Additionally, the integration of BRIDGE into *ex vivo* settings is implemented with a custom 3D-printed device that minimizes reslicing steps, enabling faster and more accurate registration in the x, y, and z axes. However, *in vivo* 2P imaging remains limited by its 600-800 µm penetration depth, restricting whole-brain coverage. The *ex vivo* pipeline offers whole brain imaging and high resolution but cannot track longitudinal changes. Therefore, bridging *in vivo* and *ex vivo* pipelines become essential for combining dynamic monitoring with high-resolution and whole brain analysis.

BRIDGE enables to develop an AI-driven MRI prediction approach using microscopic information as ground truth. Above all, it allows to use any microscopic contrast for training AI algorithms. Previously developed segmentation methodologies predominantly relying on manual or semantic annotations showed the potential of using neural networks for medical image segmentation [72, 73]. Our approach substantially augments image quality and reduces the reliance on human input, improving overall robustness and reproducibility. Without co-registered microscopy, segmentation is limited to the voxel level, as manual segmentation cannot account for hidden details within each voxel. Integrating microscopy allows deep learning to enhance segmentation beyond MRI resolution. We were able to refine and improve the resolution of MR images revealing near microscopic information of the vascular architecture in MRI. Our AI-driven analysis holds promise for training models using BRIDGE’s voxel-accurate co-registration. As these technologies evolve, BRIDGE could be translated to human tissue studies, opening new clinical opportunities for comprehensive *in vivo* imaging and diagnostics. We showed a proof-of-concept in this manuscript how to co-register human glioma tissue on MRI, paving the way for BRIDGE’s path to clinical translation.

Further, BRIDGE enables the extraction of correlative voxel intensity values, providing quantitative insights into structural changes over time. We assessed brain vessel visibility across different MRI sequences and correlated these findings with high-resolution 2P imaging. The use of a cryogenic coil enabled the detection of microvascular structures down to 20 µm, improving vessel segmentation and quantitative analyses. The distinct correlation patterns observed between TOF, T2w, and T2*w sequences with vessel diameter and blood flow velocity provide valuable insights into the strengths and limitations of each modality for vascular imaging in living brain tissue.

In the context of brain metastases, our technology platform allows characterization of the role of erythrostasis and microvascular changes in T2*w hypointense lesions. The association between T2* map positivity and reduced perimetastatic blood flow suggests that vascular dysfunction may contribute to metastatic progression. Furthermore, *ex vivo* analyses confirm increased metastatic and erythrocyte cluster formation in T2*w hypointense lesions, reinforcing the link between vascular pathology and metastasis [74]. These insights emphasize the potential of MRI-based vascular biomarkers in monitoring tumor-associated vascular changes and guiding future therapeutic strategies.

BRIDGE is able to track dynamic MR signal intensity changes of single voxels during glioblastoma progression. The distinct MRI characteristics of GG16 and S24 reflect their different growth patterns, with GG16 showing higher normalized T2w signal intensity even at similar glioma densities in 2P. Longitudinal analysis reveals that T2w variations depend on glioma density and time since tumor injection as well as tumor growth rate, subdividing hyper- and isointense glioma infiltration zone. UMAP analysis further illustrates how information from multimodal MRI aligns with glioma density measured in 2P. These findings support voxel-based MR analysis as a valuable tool for monitoring glioblastoma progression and improving imaging-based tumor assessment. Looking ahead, this approach could enable the quantification of tumor cell densities and glioma dynamics in clinical MR images.

In human glioblastoma patients, we validated that T2 signal especially in the infiltration zone not only depends on glioma cell density, but also on glioma dynamics. That is why future evaluation criteria for treatment response could include the evolving changes in T2 signal implicating glioblastoma progression [26].

In summary, BRIDGE provides a longitudinal, voxel-accurate correlation of MRI with both *in vivo* and *ex vivo* microscopy. By overcoming longstanding challenges in multimodal registration and expanding the potential of high-resolution imaging, BRIDGE stands as a powerful tool for research and will be able to affect clinical diagnostics by improving MR imaging through ground truth based deep learning.

## Abbreviations

MRI: magnetic resonance imaging
2P: two-photon microscopy
SNR: signal-to-noise-ratio
FLAIR: fluid-attenuated inversion recovery
TOF: time-of-flight imaging
T1w: T1-weighted sequence
T2w: T2-weighted sequence
T2*w: T2*-weighted
MASD: mean absolute surface distance
NGS: neuron-to-glioma synapse
TC: tumor core
IZ: infiltration zone
AI: artificial intelligence
ROI: region of interest
ExM: expansion microscopy
IHC: immunohistochemistry
VAChT: vesicular acetylcholine transporter.

## Methods

### Study participant details

Human tissues were obtained after approval of the local regulatory authorities (ethical codes 005/2003, S-672/2023, 23-1233-S1, 23-1234-S1, S-005/2003, 23-1175-S1 and PV4904).

Human patient samples were pseudonymized manually.

### Cultivation of patient-derived primary tumor cell lines and Illumina 850k methylation array characterization

Patient-derived glioblastoma cell lines were cultured from resected tumors as described in previous studies [49, 50, 64]. These cells were grown in DMEM/F-12 under serum-free, non-adherent, ‘stem-like’ conditions, supplemented with B27 (12587-010, Gibco), insulin, heparin, epidermal growth factor, and fibroblast growth factor.

Brain-tropic Jimt-1 (human breast cancer, ER-, PR-, HER2 amplification, trastuzumab-resistant, p53-/-, a kind gift from Patricia Steeg), were cultured in DMEM supplemented with 10% fetal bovine serum (FBS) and 1% penicillin/streptomycin (pen/strep) [75–77].

The molecular classification of the tumor xenograft models used in this study is available in Table S1. DNA methylation analysis of over 850,000 CpG sites in all cell lines was performed using the Illumina Infinium Methylation EPIC kit at the Genomics and Proteomics Core Facility of the German Cancer Research Center in Heidelberg, Germany [78]. Glioblastoma cell lines maintained under stem-like conditions and Jimt1 were transduced with lentiviral vectors expressing membrane-bound GFP using the pLego-T2-mGFP construct, as described before [50, 79]. The brain-tropic metastasis cell lines were stably transduced with lentiviral vectors for imaging purposes, with cytosolic expression of GFP or tdTomato achieved through transduction with plKO.1-puro-CMV-TurboGFP (SHC003, Sigma-Aldrich, USA) or cytoplasmic tdTomato (LeGo-T2, plasmid #27342, Addgene, USA). Throughout the study, all cell lines were tested for mycoplasma contamination by PCR every three months.

Regular FACS sorting of transduced cells was conducted using the FACSAria Fusion 2 (Bernhard Shoor) or FACSAria Fusion (Richard Sweet) systems. GFP-positive cells were sorted using the BL530/30 filter **(Figure S1E)**.

### Animal procedures

All animal procedures were conducted in accordance with the European Directive on animal experimentation (2010/63/EU), the Society for Laboratory animal Science (GV-SOLAS) guidelines and institutional laboratory animal research guidelines. The study was approved by the Regierungspräsidium Karlsruhe, Germany under the license numbers G50-19 and G220-16.

Male NMRI nude mice (8-12 weeks old) were used for studies involving human patient-derived primary glioblastoma cells. The S24 cell line exhibits an infiltrative growth pattern characteristic of the malignancy in human disease, while GG16 represents a tumor that is clearly visible on MRI.

Female Athymic nude mice and NSG mice (8-12 weeks old) were used for all studies involving brain metastasis models. Jimt1 cells were cultured as adherent cultures in DMEM-high glucose media supplemented with FBS and Penicillin/Streptomycin, as previously described. All cell lines were regularly sorted for fluorescence before animal procedures.

Animals were housed at the animal facilities of the German Cancer Research Center in Heidelberg.

Surgical procedures were performed according to established protocols [47, 49, 50, 64, 65]. Custom-made teflon rings instead of conventional titanium rings [15] and bespoke alignment of cross-modality *in vivo* and *ex vivo* imaging were established to ensure MRI compatibility and initial image registration **(Figure S1A)**. For MR imaging, custom-made rings crafted from Teflon were used in place of the traditional custom-made titanium ring typically employed for painless head fixation during imaging. Previous studies have reported hyperintense areas under the window caused by local edema or scar tissue shortly after cranial window surgery [16]. Our observations indicate that these artifacts regress 2-3 weeks post-surgery, making this the optimal time for correlative imaging **(Figure S1B)**.

In the xenograft glioma model, 50,000 to 100,000 tumor cells were stereotactically injected into the mouse cortex at a depth of approximately 500 µm, 1-3 weeks after cranial window surgery. Alternatively, glioma cells were injected stereotactically into the striatum without prior cranial window surgery.

For brain metastasis models, tumor cells trained in brain tropism were suspended in phosphate-buffered saline and injected into the left cardiac ventricle, following established protocols [74, 77, 80].

### Intravital microscopy

Mice that had previously undergone cranial window surgery and tumor injection were observed intravitally for up to 150 days in the glioma model and for 28 days in the brain metastases model following tumor implantation. Imaging was performed using a Zeiss 7MP microscope and a Zeiss LSM 980 with Airyscan2, both equipped with a pulsed Ti:Sapphire laser (Chameleon Discovery NX; Coherent). Fluorescent markers, including GFP, tdTomato, FITC, and TRITC dextran, were imaged using 850 nm and 960 nm wavelengths, respectively. The microscopes were configured with bandpass filter sets of 500 - 550 nm and 575 - 610 nm. A 10×, 0.45 NA and 20×, 1.0 NA, apochromatic, both with 1.7 mm working distance, water immersion objectives (Zeiss) were used for imaging on both setups. Fluorescence emission was detected using low-noise, high-sensitivity photomultiplier tubes.

Mice were anesthetized with isoflurane gas diluted in 100% O^2^. Anesthesia was induced with 3-5% isoflurane and reduced to 0.5-1.5% for maintenance during imaging, monitored by the breathing rate. Eye cream was applied to protect the eyes after anesthesia induction. Throughout the imaging process, the body temperature of the mice was maintained at 37°C using a temperature sensor and heating plate. Anesthesia depth was regularly assessed by monitoring the mice’s posture and breathing rate.

For the angiogram, TRITC-dextran (500,000 g/mol) was dissolved in a 0.9% NaCl solution at a concentration of 10 mg/ml. Before imaging, 100 µl of the TRITC or FITC solution was injected into the lateral tail vein.

Tile scans of the entire brain surface were acquired for the correlation pipeline using the 10× objective, 0.45 NA, apochromatic, water immersion, with pixel size of 1.1838 µm.

For Airyscan imaging, TRITC was used to visualize blood vessels, while the region of interest was selected based on the GFP channel, specifically highlighting the Jimt1 mGFP-expressing cells. Imaging was conducted using a 20x, 1.0 NA, apochromatic, water immersion objective (Zeiss) with a 1.7 mm working distance. The pixel size utilized was 59.54 nm, providing high-resolution imaging.

For blood flow velocity measurements, intravital imaging was performed using an intravenous angiogram with TRITC as previously described. Imaging was conducted with a 20x, 1.0 NA, apochromatic, water immersion objective (Zeiss) with a 1.7 mm working distance. A region of interest containing the vessels of interest was selected. A line scan was placed parallel to the vessel, consisting of 128 pixels with a pixel size of 1.1838 µm, resulting in a total line length of 10.6066 µm. The line was scanned repetitively for 1 second with a frame interval of 0.15 ms.

The data was processed in Zen software to generate an xy-image. Subsequent analysis was performed using a custom-written macro in Fiji, where 10 lines per line scan were drawn manually corresponding to the angle of the recorded blood flow. The mean blood flow velocity and the standard deviation were calculated.

### Intravital MRI

MRI scans were conducted using a 9.4T horizontal bore small animal MRI scanner (BioSpec 94/20 USR, Bruker BioSpin GmbH, Ettlingen, Germany) which was outfitted with a gradient strength of 675 mT/m. For S24 mice, acquisitions were conducted using an 8.4 cm body coil for transmission and a receive-only 4-channel surface array coil. Scans of mice with GG16 and Jimt1 were acquired using a cryogenic RF coil. To induce anesthesia, 4% isoflurane in 100% oxygen was used. During scanning, 1-1.5% isoflurane in 100% oxygen was administered via a nose cone. The respiratory rate was continuously monitored, and the animals were placed on a Bruker standard MRI bed with an integrated water circulation heating system to maintain body temperature. The test animals were administered a maximum of 0.1 ml of contrast agent (0.5 mmol/ml Gadolinium diluted 1:5 in PBS) intravenously. Not all of the listed sequences were scanned in every MRI session. The sequences are listed in Table S2.

### Transcardial perfusion

Brain perfusion was performed as previously described [50, 64]. Tumor-bearing mice were anesthetized using a combination of ketamine and xylazine. After confirming the absence of interphalangeal reflexes, the animals were transcardially perfused with 4% paraformaldehyde (PFA) and PBS. Following brain extraction, the tissue was fixed in 4% PFA overnight and then stored in PBS.

### 3D print device

We created a self-designed, easy-to-print and -use reference plane for reslicing by using the software FreeCAD and UltiMaker Cura and the 3D printer UltiMaker 2. The device consists of PLA filament. The main purpose of this device is to find the same slicing angle in MRI and microscopy. Besides that, the device solves the fixation of the brain to reduce MRI artifacts and gets the brain as close to the coil as possible. To use this device, the brain tissue is first cut between the midbrain and cerebellum to create a flat surface. The cut brain gets sticked on the self-designed device with superglue. Subsequently, it is pushed into a 15 ml centrifuge tube where it fits perfectly. The tube is filled with PBS and sealed so that as little air as possible remains in the tube. However, as a little air always remains in the tube, the 3D-printed device also serves as a bubble trap during MRI scans **(Figure S2A)**. To get the brain out of the tube, a simple wire was used to pull on the small opening in the device. Following, the brain is sliced on the same device to find the same reslicing angle in microscopy.

### *Ex vivo* 9.4T MRI acquisition

*Ex vivo* 9.4T MRI scans were conducted using the same scanner as described for *in vivo* scans. Scans were acquired using a cryogenic RF coil. To place the tissue as close to the coil as possible, the tissue gets sticked on the 3D print device. The sequences are listed in Table S3.

### Vibratome slicing of brain tissue

After *ex vivo* MRI acquisition, the brain tissue, which was glued onto the 3D-printed device, was removed from the 15 ml falcon tube. The 3D-printed device was then directly attached to the vibratome holding device. Brain slices, 80-100 µm thick depending on MRI voxel size, were cut using a vibratome (Leica VT1000S) to align with the voxel size of the MRI.

### Immunohistochemistry

Brain slices were either permeabilized with 5% FBS and 1% Triton X-100 (TX100) for 2 hours at room temperature or underwent antigen retrieval using 20 mM sodium citrate, incubated at 60°C for 1 hour. After antigen retrieval, the slices were washed with 10% FBS.

Primary antibodies were incubated in 1% FBS and 0.2% TX100 (anti-GFP chicken, Abcam ab13970, 1:1000; anti-RFP guinea pig, SySy REF#390004, 1:200; anti-Ter119 rat, Thermo Fisher Invitrogen, REF#14-5921-85, 1:500) overnight on a shaking device at 4°C. Slices were then washed twice in 2% FBS, followed by incubation with the secondary antibody at a 1:500 dilution in 1% FBS and 0.2% TX100 for at least 3 hours at room temperature. After secondary antibody incubation, slices were washed three times with 1% FBS and then three times with PBS. Finally, slices were washed with 1:10000 DAPI in PBS before widefield microscopy. Slices were mounted on microscope slides with SlowFade gold.

### Human immunohistochemistry

Brain slices either underwent antigen retrieval by heating 20 mM sodium citrate buffer in a microwave until boiling, immediately transferring the hot buffer to the wells containing the tissue and then incubated at 60°C for 30 minutes or permeabilized with 5% FBS and 1% Triton X-100 (TX100) for 2 hours at room temperature. Following antigen retrieval, slices were blocked in 5% FBS in PBS for 2 hours at room temperature.

Primary antibody incubation was performed overnight at 4°C on a shaker in a solution containing 1% FBS and 0.2% Triton X-100 in PBS (anti-Nestin, mouse, 1:300, Abcam, ab22035, anti-CD31, goat, R&D systems, AF3628, 1:100). After incubation, slices were washed three times for 10-15 minutes each with 2% FBS in PBS.

Secondary antibodies (Alexa Fluor 488 donkey anti-mouse for Nestin; Alexa Fluor 647 donkey anti-rat for CD31) were diluted 1:500 in 1% FBS in PBS (without Triton X-100) and incubated either overnight at 4°C or for 3 hours at room temperature on a shaker. After secondary antibody incubation, slices were washed three times for 10–15 minutes each in 1% FBS in PBS, followed by a final 10–15 minute wash with PBS alone.

For expansion microscopy, only Nestin-stained slices were used, and the Alexa Fluor 488-conjugated secondary antibody was replaced by Alexa Fluor 568. Finally, all slices were mounted with SlowFade Gold containing DAPI and imaged using widefield fluorescence microscopy.

### Widefield microscopy of *ex vivo* slices

Widefield microscopes (Leica DM6000, Leica Mica, Zeiss AxioScanZ1) were used to acquire multichannel tile scans of stained *ex vivo* slices. For quantitative analyses, the brain slices were imaged at the AxioScanZ1 microscope with a pixel size of 325 nm using a 20x (NA 0.8) objective.

### Expansion microscopy

Brain slices were prepared and stained as described in the immunohistochemistry section, with the exception of using Atto 647N instead of Alexa 647, and staining endogenous mGFP with anti-GFP as described previously. Both primary and secondary antibody incubation times were extended to 24 hours. Following staining, slices were screened using a widefield fluorescence microscope (Leica DM6000), and small regions of interest, approximately 1 cm x 1 cm in size, were cut out using a scalpel.

For sample anchoring, the trimmed tissues were incubated in 0.1 mg/ml Acryloyl-X SE (AcX) solution diluted in 1x PBS overnight at room temperature without shaking. Right before the gelation procedure, a gelation chamber was prepared by placing two 1.5 coverslips, cut with a diamond knife, on top of each other at the edges of a parafilm-coated objective slide. The samples were then placed onto a Poly-L-Lysin-coated objective slide.

To prepare the gelation solution, 470 μl of monomer solution (composed of 0.08% [v/v] sodium acrylate [33% wt stock], 2.5% [v/v] acrylamide [50% wt stock], 0.02% [v/v] cross-linker [1% wt stock], 1.9M NaCl [5M stock], 1 ml of 10× PBS, and 18.8% [v/v] water) was combined with 10 μl each of 0.5 wt% 4-HT, 10 wt% TEMED, and 10 wt% APS. All stock solutions were prepared with MilliQ water. Then, 60 μl of the freshly prepared gelation solution was applied between the spacers, and the Poly-L-Lysin slide with the tissue was placed on top of the gelation solution. Radical polymerization was initiated by incubating the gelation chamber at 37°C for 2 hours.

Post-polymerization, the gel was trimmed into a trapezoid shape to distinguish the orientation of the sample and transferred into a 12-well plate containing 1 ml of digestion buffer (for 50 ml: 250 μl Triton-X100, 100 μl EDTA, 2.5 ml Tris-Cl, 2.338 g NaCl, filled up with MilliQ water – with 1:100 Proteinase K [800U/ml] freshly added before the experiment). The samples were incubated overnight at room temperature on a shaker.

The next day, the gel was washed twice with 1x PBS for 30 minutes each. Subsequently, the gel was transferred into a 6-well plate with MilliQ water. The water was exchanged every 10 minutes for three cycles, followed by an additional 30-minute wash. The samples were then trimmed and mounted on Poly-L-Lysin-coated glass-bottom dishes (MatTek) for imaging, and were covered with MilliQ water.

For Airyscan microscopy, the expanded sample was imaged using an LSM900 Airyscan NIR microscope (Zeiss) equipped with a 40x, 1.2 NA water immersion objective. Images were acquired with a resolution of 59.54 nm. Airyscan post-processing was performed using Zen Blue software.

### Electron microscopy

The procedure for sample preparation was carried out in accordance with previously established methods [77]. Two weeks following intracardial injection, the animals bearing brain metastases were administered a deep anesthesia using a combination of ketamine and xylazine. Subsequently, they were perfused transcardially with a solution containing 4% (w/v) paraformaldehyde (PFA) in 1x phosphate-buffered saline (PBS, Sigma). Upon removal, the brains were postfixed in a 4% PFA solution for a duration of four hours.

Sections of 100-200 μm thickness were then cut using a vibratome (Leica VT1000S). The slices from the xenograft brain were examined under a widefield fluorescence microscope (Leica DM6000) to detect the endogenous fluorescence of brain metastatic cells. For DAB labeling, samples were next immersed in a 10% (w/v) sucrose solution (Sigma) in PBS for a duration of 10 minutes, followed by a 12-15 hour incubation in a 30% sucrose solution. They were then subjected to a freeze-thaw process in liquid nitrogen twice, each for 5 seconds, before being placed in a blocking solution composed of 5% FBS in PBS, and left at room temperature (RT) for 1 hour. The slices were incubated overnight at 4°C with specific antibodies. Alternatively, the samples were processed after dissection at the widefield microscope with a heavy metal stain without DAB labeling and metastatic cells were ultrastructurally identified.

For the DAB precipitate labeling, samples were incubated with the secondary antibody, a biotinylated anti-mouse AB (abcam (ab6788), 1:500, in blocking solution), for 12-15 hours at 4°C. After washing thrice with PBS, the samples were treated with Vectastain ABC-kit (Linaris) solution for 1 hour at RT. The samples were then incubated in a glucose-DAB solution (glucose: 2 mg/ml, DAB: 1.4 mg/ml, dissolved in PBS) for 10 minutes, followed by a 1-hour incubation in a glucose-DAB-glucose oxidase solution (glucose oxidase: 0.1 mg/ml, Serva). This process facilitated the formation of an electron-dense precipitate. The success of the reaction was monitored using widefield light microscopy. All samples were subsequently processed as described previously, embedded in resin and cut with an ultramicrotome (Ultracut S, Leica).

EM images were acquired using an Aquilos system with Maps Viewer 3.1.0.

### Image deconvolution

For content-aware restoration, training data was collected following established methodologies [50] and trained using CSBDeep [25]. The prediction outputs were utilized as prediction maps in Ilastik [39]. Subsequently, the raw data was segmented using Ilastik Autocontext. Registered probability maps were then binarized to values 0 and 1 using a threshold of 10000 in Fiji and converted to 32-bit. 2P stacks were grouped in z axis by average to match MRI voxel depth so that one single 2P voxel depicts the glioma volume density from 0 to 1.

### *In vivo* registration workflow

The registration workflow consists of the following steps: (1) resampling of 2P z-slices by grouping them to match the voxel depth of the MRI, (2) orientation alignment of both 2P and MR images using the cranial window glass as a common reslicing plane, (3) spatial resolution resampling of MRI in xy to match 2P pixel size, and (4) elastic, landmark-based registration, enabling precise voxel-to-voxel correlation.

To be more specific, both MRI and 2P data underwent manual reslicing in Fiji. The Reslice function was used to align the data parallel to the cranial window, ensuring that both MRI and 2P images shared the same 3D orientation which is of particular importance for our workflow. The MRI data were then upscaled to match the resolution of the 2P data in the x and y axes. Subsequently, elastic registration was performed using the thin-plate mode of the “Landmark registration” module in 3D Slicer. Affine and rigid transformations were implemented following standard methodologies [81] while elastic registration was performed using free-form deformations [82]. Landmark-based registration was guided by the principles of thin-plate splines [83]. The MR image was used as fixed volume, the 2P image as moving volume. T2w images were used for registrations of the glioma cohorts, T2* maps were used for registrations of the metastases cohorts. Manual landmark registration utilizing anatomical landmarks of blood vessels identifiable in both MRI and 2P imaging enables a precise registration of both vessel and tumor channels to match MRI. For local refinement, the Simple ITK option was selected. Final transformation was conducted using the 3D Slicer module “Resample Scalar/Vector/DWI Volume” by providing the transformation file resulting from the previous module, the MRI image as fixed volume and the 2P image as moving volume.

### MRI image processing

To ensure consistency of weighted imaging sequences across datasets for *in vivo* 2P-MRI correlation, MR normalization was conducted using a normal appearing gray matter z-normalization as only gray matter was analyzed [84] **(Figure S1C)**. Therefore, custom masks of every registered dataset were created based on 2P signal excluding tumor signal, vessels, lesion artifact due to injection and other artifacts and dividing the mask into two categories: normal appearing gray matter (category ‘w/o glioma’ in Figure 6) and glioma (other categories in Figure 6). These masks were also used to determine voxels for quantification. Mapping sequences were fitted using the Fiji plugin FijiRelax [85]. Different sequences were registered using a custom-code registration Fiji macro relying on rotation and translation.

Clinical MR images were z-normalized to cerebrospinal fluid (CSF).

### Voxel intensity extraction

To enable a voxel-by-voxel correlative quantification, x and y axes of 2P were downscaled by bilinear interpolation to match the MRI resolution **(Figure S5A)**. The resulting downscaled 2P image consists of values from 0 to 1 representing the glioma density within this voxel. Using a self written Fiji macro, voxel intensities of the normalized MR and 2P images were extracted inside the mask described above.

### Longitudinal registration and quantification of tumor growth

After registration of each dataset on its own, the datasets of each mouse were registered to the first time point using a self-written Fiji macro relying on rotation and translation. This step enabled defining MRI and 2P value changes per day. Masks for quantification of glioma growth were created by overlapping the masks of every timepoint per mouse.

### RMSE quantification for sequence comparison

Root mean square error (RMSE) for sequence comparison was detected by defining 5 corresponding landmarks in MRI and 2P microscopy after registration. RMSE was calculated as follows:

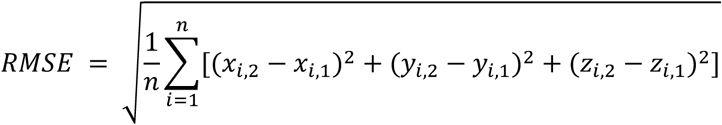

### Inter-rater reliability

Two raters familiar with the workflow registered the same six 2P microscopy datasets to already resliced T2w images. To determine interrater reliability, a coordinate system consisting of 50 columns and 50 rows per slice were applied to the grouped, but not resliced 2P images. Using this coordinate system which gets transformed during reslicing and registration, we were able to define the average RMSE per dataset between the same randomly suggested coordinates of both rater’s registrations with a self-written Fiji macro.

### *Ex vivo* registration workflow

Similar to the *in vivo* registration pipeline, MRI scans were resliced in Fiji using the 3D-printed device as a reslicing plane. The x and y axes of the MRI images were upscaled to match the resolution of the microscopy images. There was no need to reslice the microscopy images, as they were already captured with the correct reslice angle due to the use of the 3D-printed device. Subsequently, the corresponding MRI slices were extracted that matched the widefield image slice stack. Since the voxel depth of the MRI and the slicing depth of the brain sections were identical, the subsequent slices were automatically aligned.

For the next step of elastic registration, the MRI images were used as fixed volumes, and the microscopy images as moving volumes. The ThinPlate mode of the “Landmark Registration” module in 3D Slicer was utilized for this process. For local refinement, the Simple ITK option was selected. The DAPI channel of the widefield images was used for registration to identify anatomical landmarks such as the corpus callosum or the anterior commissure. In most cases, a few key landmarks at prominent points were sufficient to achieve perfect registration between the modalities.

The final transformation was conducted using the 3D Slicer module “Resample Scalar/Vector/DWI Volume” by providing the transformation file generated from the previous module, with the MRI image as the fixed volume and the microscopy image as the moving volume. Landmark registration was then applied to the other channels of the widefield images. All registered widefield images were subsequently processed using Ilastik pixel classification, generating probability maps as the output.

### Semi-automatic quantification of brain metastases

*Ex vivo* Jimt1 tdtomato brain slices stained with anti-RFP, anti-Ter119 and DAP were registered onto T2*w MRI and probability maps were trained in Ilastik. Manual ROIs were drawn and labeled according to specific metastases. Images were thresholded and the function Analyze Particles was applied to the ROIs to measure metastatic cell clusters and erythrocyte clusters in custom written Fiji macros. The resulting measurement tables were subsequently merged and analyzed in R.

### *In vivo*-*ex vivo* registration workflow

*In vivo* and *ex vivo* MRI were elastically registered in 3D Slicer using the *ex vivo* MRI as the fixed volume. Thereby, *in vivo* MRI is automatically registered onto light microscopy data using *ex vivo* MRI for bridging between light microscopy and *in vivo* MRI.

### *Ex vivo* atlas registration

To register brain sections onto the Allen Mouse Brain Common Coordinate Framework (CCFv3) [46], we used the QUINT workflow [86] **(Figure S2C)**.

### Deep learning workflow in BRIDGE

The datasets, registered using the BRIDGE pipeline, were resampled to achieve uniform voxel spacing in the XY plane with a voxel size of 25 µm, while maintaining a voxel size of 100 µm in the Z direction. We established a minimum vessel diameter filter at 30 µm for 2P data as the approximate visibility threshold for vessels in the T2w sequence **(Figure 5F)**. This protected the neuronal network from a possible confusion caused by many small vessels **(Figure 4A)**.

Small vessels less than 30 μm in diameter were excluded to avoid confusion of the network. Therefore, a custom Fiji macro was developed to filter out structures with diameters below 30 µm. The macro systematically analyzed the local neighborhood of each voxel to estimate the vessel width. By retaining only vessels meeting the size threshold, the filtering process enhanced the relevance of the data for subsequent analysis and model training. Single slice crops were manually selected from regions with high registration accuracy and clear vessel visibility, ensuring the inclusion of areas with optimal image quality and well-defined vasculature. Vessel signal was binarized to create vessel segmentations. To increase data diversity and improve model generalization, random flips and rotations were applied as data augmentation. These transformations generated additional variations of the original data, reducing overfitting risks and enhancing the robustness of the trained model. We used the 2D training workflow in nnU-Net [24] to train the MR images with the preprocessed ground truth. To assess the training, we performed a 5-fold cross-validation **(Figure S3A)**. We used the Dilated Dice coefficient [51] since the structures we evaluated were small and thin. In the calculation, the prediction is compared to the registered microscopy ground truth, so the Dice coefficient would also include the registration error. To compensate for the registration error, we used the Dilated Dice coefficient instead of the Dice coefficient. For dilation, 50 µm were added on both sides of the vessels. The Dilated Dice coefficient was calculated as follows:

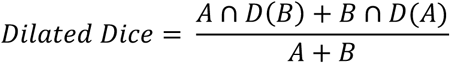

A is the microscopy ground truth, B is the prediction, D(A) is the dilation of the microscopy ground truth and D(B) is the dilation of the prediction.

Further, the mean absolute surface distance (MASD) [52] was calculated as follows:

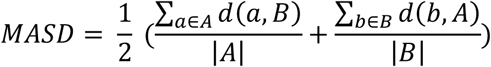

A is the microscopy ground truth, B is the prediction. a is a voxel in the microscopy ground truth, b is a voxel in the prediction. d(a,B) is the distance of a voxel in the ground truth to the closest voxel of the prediction. d(b,A) is the distance of a voxel in the prediction to the closest voxel of the ground truth. |A| is the number of ground truth voxels. |B| is the number of prediction voxels.

Additionally, all vessels in the MR image and the prediction were classified into three categories: “strongly visible,” “barely visible,” and “not visible.” Their diameters were also compared to the ground truth. Vessel diameters in MRI were determined by thresholding the MR image.

### Statistical analyses

Plots were generated with the ggplot2 package and statistically tested with ggpubr from R with RStudio. Boxplots indicate minimum and maximum, median, and the first and third quartiles. For data visualization in Plots, further packages were used: gridExtra, umap, viridis, fields, RColorBrewer, plotly and cowplot.

* = P<0.05; ** = P<0.01, *** = P<0.001

### 3D-Visualization

3D image stacks were loaded into arivis 4D. Pixel sizes and contrast thresholds were adjusted. For visualizing MRI in 3D, skull-stripping was performed using a custom trained Unet-model. High resolution images were then taken in the ZEISS arivis software.

### Schematic illustrations

Schematics of experimental workflows were created with BioRender.com.

### Use of LLM

To assist coding and to improve the manuscript’s readability, the authors utilized ChatGPT-4o and ChatGPT-5. Following its use, the authors thoroughly reviewed and made any necessary edits to the content. The authors assume full responsibility for the final results.

## Data availability

Source data are provided with this paper. Raw data are available on reasonable request.

## Code availability

Custom codes can be found on https://github.com/venkataramani-lab.

## Acknowledgements

V.V. and M.K. received financial support from the European Center for Neuro-Oncology (EZN), German Research Foundation (DFG: VE1373/2-1), the Else Kröner-Fresenius-Stiftung (2020-EKEA.135), the Medical Faculty of Heidelberg University (Physician-Scientist-Program, Krebs-und Scharlachstiftung) and SFB 1389, UNITE Glioblstoma, project ID 404521405 (addressed to V.V., E.R., M.A.K.). Y.Y., J.S., and E.R. were supported by the Deutsche Krebshilfe/German Cancer Aid (Mildred-Scheel-Scholarship for MD students). V.V. and M.A.K. were supported by a grant from the German Research Foundation (DFG: SFB 1389). M.A.K. received financial support from the Bundesministerium für Bildung und Forschung (BMBF) within the framework of the medical research and funding concept (01ZX1913D) and from the DFG (Project number 259332240/RTG 2099). M.A.K. was also supported by a grant from the Hertie Foundation for Excellence in Clinical Neuroscience. V.V. was supported by the Schwiete-Stiftung and the Wilhelm-Sander-Stiftung. We gratefully acknowledge the data storage service SDS@hd, supported by the Ministry of Science, Research, and the Arts Baden-Württemberg (MWK). O.T.H. received financial support from the Heidelberg University Medical Faculty Clinician Scientist Program.

We thank S. Kegel and M. Ladd for the support and opportunity to use the 9.4T MRI scanner at the DKFZ. We thank Laura Fankhauser for her help and introduction to 2P and MRI infrastructures for the initial experiments. We thank Nina Drewa for her help with cell culture and transductions. We thank Yvette Dörflinger, Simone Hoppe, and Niklas Wißmann for their assistance with sample preparation and electron microscopy image analysis. We acknowledge Jovana Bojcevski and Amir Abdollahi for providing xenograft brain tissue, and Hannah Fels-Palesandro for her help with *ex vivo* MRI. We thank Volker Sturm for his assistance with MRI sequence alignment. We thank Gerald Bendner for his advice on antigen retrieval.

We gratefully acknowledge the support of Manuela Schulz, Damir Krunic, and Felix Bestvater from the Light Microscopy Core Facility of the German Cancer Research Center. We thank F. Blum, T. Rubner, M. Eich, K. Hexel, and S. Schmitt from the Flow Cytometry Core Facility for their help with FACS experiments. We thank R. Matejka from the precision mechanics department of the German Cancer Research center for his assistance in designing and manufacturing of the teflon head fixation device. Finally, we thank K. Dell, A. Riedasch, K. Schmidt, and A. Berdel for their support with animal care and the design of animal experiments.

We acknowledge the use of AI tools to assist with coding and to improve the manuscript’s readability.

## Author information

These authors contributed equally: Julian Schroers, Yvonne Yang.

## Contributions

Supervision, V.V.; conceptualization, J.S., Y.Y., and V.V.; methodology, J.S., Y.Y., E.R., E.I.Z., project administration: J.S., Y.Y., and V.V.; investigation, J.S., Y.Y., E.R., N.S., E.I.Z., A.H., J.G.S., A.P.L., O.T.A., T.M., M.F.; formal analysis J.S., Y.Y., E.R., E.I.Z., A.H. and V.V.; resources, J.J., M.K., F.L.R., F.K. and V.V. visualization, J.S., Y.Y, V.V., ; writing original draft J.S. and Y.Y.; writing – review and editing, J.S., Y.Y., O.T.A., J.J., B.S., D.H.H., M.K., M.O.B., F.K. and V.V.; funding acquisition, F.K., V.V.

## References

1. Kabasawa H. MR Imaging in the 21st Century: Technical Innovation over the First Two Decades. Magn Reson Med Sci. 2022; 21: 71–82.

2. Langen KJ, Galldiks N, Hattingen E, Shah NJ. Advances in neuro-oncology imaging. Nat Rev Neurol. 2017; 13: 279–89.

3. Lemee JM, Clavreul A, Menei P. Intratumoral heterogeneity in glioblastoma: don’t forget the peritumoral brain zone. Neuro Oncol. 2015; 17: 1322–32.

4. Wen PY, van den Bent M, Youssef G, Cloughesy TF, Ellingson BM, Weller M, et al. RANO 2.0: Update to the Response Assessment in Neuro-Oncology Criteria for High- and Low-Grade Gliomas in Adults. J Clin Oncol. 2023; 41: 5187–99.

5. Weinberg BD, Gore A, Shu H-KG, Olson JJ, Duszak R, Voloschin AD, et al. Management-Based Structured Reporting of Posttreatment Glioma Response With the Brain&#xa0;Tumor Reporting and Data System. Journal of the American College of Radiology. 2018; 15: 767–71.

6. Parillo M, Mallio CA, Pileri M, Dirawe D, Romano A, Bozzao A, et al. Interrater reliability of Brain Tumor Reporting and Data System (BT-RADS) in the follow up of adult primary brain tumors: a single institution experience in Italy. Quantitative Imaging in Medicine and Surgery. 2023; 13: 7423–31.

7. Wen PY, Macdonald DR, Reardon DA, Cloughesy TF, Sorensen AG, Galanis E, et al. Updated Response Assessment Criteria for High-Grade Gliomas: Response Assessment in Neuro-Oncology Working Group. Journal of Clinical Oncology. 2010; 28: 1963–72.

8. Gillies RJ, Kinahan PE, Hricak H. Radiomics: Images Are More than Pictures, They Are Data. Radiology. 2016; 278: 563–77.

9. Aerts HJWL, Velazquez ER, Leijenaar RTH, Parmar C, Grossmann P, Carvalho S, et al. Decoding tumour phenotype by noninvasive imaging using a quantitative radiomics approach. Nature Communications. 2014; 5: 4006.

10. Lambin P, Rios-Velazquez E, Leijenaar R, Carvalho S, van Stiphout RGPM, Granton P, et al. Radiomics: Extracting more information from medical images using advanced feature analysis. European Journal of Cancer. 2012; 48: 441–6.

11. Brugnara G, Baumgartner M, Scholze ED, Deike-Hofmann K, Kades K, Scherer J, et al. Deep-learning based detection of vessel occlusions on CT-angiography in patients with suspected acute ischemic stroke. Nature Communications. 2023; 14: 4938.

12. Rastogi A, Brugnara G, Foltyn-Dumitru M, Mahmutoglu MA, Preetha CJ, Kobler E, et al. Deep-learning-based reconstruction of undersampled MRI to reduce scan times: a multicentre, retrospective, cohort study. Lancet Oncol. 2024; 25: 400–10.

13. Wang X, Leong ATL, Tan SZK, Wong EC, Liu Y, Lim L-W, et al. Functional MRI reveals brain-wide actions of thalamically-initiated oscillatory activities on associative memory consolidation. Nature Communications. 2023; 14: 2195.

14. Desjardins M, Kılıç K, Thunemann M, Mateo C, Holland D, Ferri CGL, et al. Awake Mouse Imaging: From Two-Photon Microscopy to Blood Oxygen Level–Dependent Functional Magnetic Resonance Imaging. Biological Psychiatry: Cognitive Neuroscience and Neuroimaging. 2019; 4: 533–42.

15. Karimian-Jazi K, Münch P, Alexander A, Fischer M, Pfleiderer K, Piechutta M, et al. Monitoring innate immune cell dynamics in the glioma microenvironment by magnetic resonance imaging and multiphoton microscopy (MR-MPM). Theranostics. 2020; 10: 1873–83.

16. Asan L, Falfán-Melgoza C, Beretta CA, Sack M, Zheng L, Weber-Fahr W, et al. Cellular correlates of gray matter volume changes in magnetic resonance morphometry identified by two-photon microscopy. Scientific Reports. 2021; 11: 4234.

17. Bhargava A, Monteagudo B, Kushwaha P, Senarathna J, Ren Y, Riddle RC, et al. VascuViz: a multimodality and multiscale imaging and visualization pipeline for vascular systems biology. Nature Methods. 2022; 19: 242–54.

18. Lake EMR, Ge X, Shen X, Herman P, Hyder F, Cardin JA, et al. Simultaneous cortex-wide fluorescence Ca2+ imaging and whole-brain fMRI. Nature Methods. 2020; 17: 1262–71.

19. Mandino F, Horien C, Shen X, Desrosiers-Gregoire G, Luo W, Markicevic M, et al. Multimodal identification of the mouse brain using simultaneous Ca (2+) imaging and fMRI. bioRxiv. 2025.

20. Vafaii H, Mandino F, Desrosiers-Gregoire G, O’Connor D, Markicevic M, Shen X, et al. Multimodal measures of spontaneous brain activity reveal both common and divergent patterns of cortical functional organization. Nat Commun. 2024; 15: 229.

21. Johnson GA, Tian Y, Ashbrook DG, Cofer GP, Cook JJ, Gee JC, et al. Merged magnetic resonance and light sheet microscopy of the whole mouse brain. Proc Natl Acad Sci U S A. 2023; 120: e2218617120.

22. Schregel K, Heinz L, Hunger J, Pan C, Bode J, Fischer M, et al. A Cellular Ground Truth to Develop MRI Signatures in Glioma Models by Correlative Light Sheet Microscopy and Atlas-Based Coregistration. The Journal of Neuroscience. 2023; 43: 5574–87.

23. Breckwoldt MO, Bode J, Sahm F, Krüwel T, Solecki G, Hahn A, et al. Correlated MRI and Ultramicroscopy (MR-UM) of Brain Tumors Reveals Vast Heterogeneity of Tumor Infiltration and Neoangiogenesis in Preclinical Models and Human Disease. Front Neurosci. 2018; 12: 1004.

24. Isensee F, Jaeger PF, Kohl SAA, Petersen J, Maier-Hein KH. nnU-Net: a self-configuring method for deep learning-based biomedical image segmentation. Nature Methods. 2021; 18: 203–11.

25. Weigert M, Schmidt U, Boothe T, Muller A, Dibrov A, Jain A, et al. Content-aware image restoration: pushing the limits of fluorescence microscopy. Nat Methods. 2018; 15: 1090–7.

26. Li M, Huang W, Chen H, Jiang H, Yang C, Shen S, et al. T2/FLAIR Abnormity Could be the Sign of Glioblastoma Dissemination. Front Neurol. 2022; 13: 819216.

27. Radbruch A, Lutz K, Wiestler B, Baumer P, Heiland S, Wick W, et al. Relevance of T2 signal changes in the assessment of progression of glioblastoma according to the Response Assessment in Neurooncology criteria. Neuro Oncol. 2012; 14: 222–9.

28. Artzi M, Bokstein F, Blumenthal DT, Aizenstein O, Liberman G, Corn BW, et al. Differentiation between vasogenic-edema versus tumor-infiltrative area in patients with glioblastoma during bevacizumab therapy: a longitudinal MRI study. Eur J Radiol. 2014; 83: 1250–6.

29. Lutz K, Wiestler B, Graf M, Bäumer P, Floca R, Schlemmer HP, et al. Infiltrative patterns of glioblastoma: Identification of tumor progress using apparent diffusion coefficient histograms. J Magn Reson Imaging. 2014; 39: 1096–103.

30. Upadhyay N, Waldman AD. Conventional MRI evaluation of gliomas. Br J Radiol. 2011; 84 Spec No 2: S107-11.

31. Chen F, Tillberg PW, Boyden ES. Expansion microscopy. Science. 2015; 347: 543–8.

32. Wassie AT, Zhao Y, Boyden ES. Expansion microscopy: principles and uses in biological research. Nat Methods. 2019; 16: 33–41.

33. Klimas A, Gallagher BR, Wijesekara P, Fekir S, DiBernardo EF, Cheng Z, et al. Magnify is a universal molecular anchoring strategy for expansion microscopy. Nature Biotechnology. 2023; 41: 858–69.

34. Zhao Y, Bucur O, Irshad H, Chen F, Weins A, Stancu AL, et al. Nanoscale imaging of clinical specimens using pathology-optimized expansion microscopy. Nature Biotechnology. 2017; 35: 757–64.

35. Tillberg PW, Chen F, Piatkevich KD, Zhao Y, Yu C-C, English BP, et al. Protein-retention expansion microscopy of cells and tissues labeled using standard fluorescent proteins and antibodies. Nature Biotechnology. 2016; 34: 987–92.

36. Chozinski TJ, Halpern AR, Okawa H, Kim H-J, Tremel GJ, Wong ROL, et al. Expansion microscopy with conventional antibodies and fluorescent proteins. Nature Methods. 2016; 13: 485–8.

37. Navarro KL, Huss M, Smith JC, Sharp P, Marx JO, Pacharinsak C. Mouse Anesthesia: The Art and Science. ILAR Journal. 2021; 62: 238–73.

38. Tu Z, Bai X. Auto-Context and Its Application to High-Level Vision Tasks and 3D Brain Image Segmentation. IEEE Transactions on Pattern Analysis and Machine Intelligence. 2010; 32: 1744–57.

39. Berg S, Kutra D, Kroeger T, Straehle CN, Kausler BX, Haubold C, et al. ilastik: interactive machine learning for (bio)image analysis. Nature Methods. 2019; 16: 1226–32.

40. Haacke EM, Xu Y, Cheng Y-CN, Reichenbach JR. Susceptibility weighted imaging (SWI). Magnetic Resonance in Medicine. 2004; 52: 612–8.

41. Reichenbach JR, Venkatesan R, Schillinger DJ, Kido DK, Haacke EM. Small vessels in the human brain: MR venography with deoxyhemoglobin as an intrinsic contrast agent. Radiology. 1997; 204: 272–7.

42. Huff J. The Airyscan detector from ZEISS: confocal imaging with improved signal-to-noise ratio and super-resolution. Nature Methods. 2015; 12: i–ii.

43. Wu X, Hammer JA. ZEISS Airyscan: Optimizing Usage for Fast, Gentle, Super-Resolution Imaging. Methods Mol Biol. 2021; 2304: 111–30.

44. Scheffler M, Maturana E, Salomir R, Haller S, Kövari E. Air bubble artifact reduction in post-mortem whole-brain MRI: the influence of receiver bandwidth. Neuroradiology. 2018; 60: 1089–92.

45. Kwok WE. Basic Principles of and Practical Guide to Clinical MRI Radiofrequency Coils. RadioGraphics. 2022; 42: 898–918.

46. Wang Q, Ding SL, Li Y, Royall J, Feng D, Lesnar P, et al. The Allen Mouse Brain Common Coordinate Framework: A 3D Reference Atlas. Cell. 2020; 181: 936–53 e20.

47. Tetzlaff SK, Reyhan E, Layer N, Bengtson CP, Heuer A, Schroers J, et al. Characterizing and targeting glioblastoma neuron-tumor networks with retrograde tracing. Cell. 2025; 188: 390–411.e36.

48. Venkataramani V, Schneider M, Giordano FA, Kuner T, Wick W, Herrlinger U, et al. Disconnecting multicellular networks in brain tumours. Nat Rev Cancer. 2022; 22: 481–91.

49. Venkataramani V, Tanev DI, Strahle C, Studier-Fischer A, Fankhauser L, Kessler T, et al. Glutamatergic synaptic input to glioma cells drives brain tumour progression. Nature. 2019; 573: 532–8.

50. Venkataramani V, Yang Y, Schubert MC, Reyhan E, Tetzlaff SK, Wißmann N, et al. Glioblastoma hijacks neuronal mechanisms for brain invasion. Cell. 2022; 185: 2899–917.e31.

51. Trutti AC, Fontanesi L, Mulder MJ, Bazin PL, Hommel B, Forstmann BU. A probabilistic atlas of the human ventral tegmental area (VTA) based on 7 Tesla MRI data. Brain Struct Funct. 2021; 226: 1155–67.

52. Benes M, Zitova B. Performance evaluation of image segmentation algorithms on microscopic image data. J Microsc. 2015; 257: 65–85.

53. Wardlaw JM, Smith EE, Biessels GJ, Cordonnier C, Fazekas F, Frayne R, et al. Neuroimaging standards for research into small vessel disease and its contribution to ageing and neurodegeneration. Lancet Neurol. 2013; 12: 822–38.

54. Zhang L, Zhu Y, Qi Y, Wan L, Ren L, Zhu Y, et al. T(2)-Weighted Whole-Brain Intracranial Vessel Wall Imaging at 3 Tesla With Cerebrospinal Fluid Suppression. Front Neurosci. 2021; 15: 665076.

55. MacDonald ME, Frayne R. Cerebrovascular MRI: a review of state-of-the-art approaches, methods and techniques. NMR Biomed. 2015; 28: 767–91.

56. Breckwoldt MO, Bode J, Kurz FT, Hoffmann A, Ochs K, Ott M, et al. Correlated magnetic resonance imaging and ultramicroscopy (MR-UM) is a tool kit to assess the dynamics of glioma angiogenesis. eLife. 2016; 5: e11712.

57. Parker DL, Yuan C, Blatter DD. MR angiography by multiple thin slab 3D acquisition. Magn Reson Med. 1991; 17: 434–51.

58. Bollmann S, Mattern H, Bernier M, Robinson SD, Park D, Speck O, et al. Imaging of the pial arterial vasculature of the human brain in vivo using high-resolution 7T time-of-flight angiography. eLife. 2022; 11: e71186.

59. Baltes C, Radzwill N, Bosshard S, Marek D, Rudin M. Micro MRI of the mouse brain using a novel 400 MHz cryogenic quadrature RF probe. NMR Biomed. 2009; 22: 834–42.

60. Franceschi AM, Moschos SJ, Anders CK, Glaubiger S, Collichio FA, Lee CB, et al. Use of Susceptibility-Weighted Imaging (SWI) in the Detection of Brain Hemorrhagic Metastases from Breast Cancer and Melanoma. J Comput Assist Tomogr. 2016; 40: 803–5.

61. Namdee K, Carrasco-Teja M, Fish MB, Charoenphol P, Eniola-Adefeso O. Effect of Variation in hemorheology between human and animal blood on the binding efficacy of vascular-targeted carriers. Scientific Reports. 2015; 5: 11631.

62. Kina T, Ikuta K, Takayama E, Wada K, Majumdar AS, Weissman IL, et al. The monoclonal antibody TER-119 recognizes a molecule associated with glycophorin A and specifically marks the late stages of murine erythroid lineage. Br J Haematol. 2000; 109: 280–7.

63. Joseph JV, van Roosmalen IA, Busschers E, Tomar T, Conroy S, Eggens-Meijer E, et al. Serum-Induced Differentiation of Glioblastoma Neurospheres Leads to Enhanced Migration/Invasion Capacity That Is Associated with Increased MMP9. PLoS One. 2015; 10: e0145393.

64. Osswald M, Jung E, Sahm F, Solecki G, Venkataramani V, Blaes J, et al. Brain tumour cells interconnect to a functional and resistant network. Nature. 2015; 528: 93–8.

65. Schubert MC, Soyka SJ, Tamimi A, Maus E, Schroers J, Wißmann N, et al. Deep intravital brain tumor imaging enabled by tailored three-photon microscopy and analysis. Nature Communications. 2024; 15: 7383.

66. Dasgupta A, Geraghty B, Maralani PJ, Malik N, Sandhu M, Detsky J, et al. Quantitative mapping of individual voxels in the peritumoral region of IDH-wildtype glioblastoma to distinguish between tumor infiltration and edema. J Neurooncol. 2021; 153: 251–61.

67. He L, Zhang H, Li T, Yang J, Zhou Y, Wang J, et al. Distinguishing Tumor Cell Infiltration and Vasogenic Edema in the Peritumoral Region of Glioblastoma at the Voxel Level via Conventional MRI Sequences. Academic Radiology. 2024; 31: 1082–90.

68. Becht E, McInnes L, Healy J, Dutertre C-A, Kwok IWH, Ng LG, et al. Dimensionality reduction for visualizing single-cell data using UMAP. Nature Biotechnology. 2019; 37: 38–44.

69. Oltmer J, Slepneva N, Llamas Rodriguez J, Greve DN, Williams EM, Wang R, et al. Quantitative and histologically validated measures of the entorhinal subfields in ex vivo MRI. Brain Communications. 2022; 4: fcac074.

70. Scarpelli ML, Healey DR, Mehta S, Kodibagkar VD, Quarles CC. A practical method for multimodal registration and assessment of whole-brain disease burden using PET, MRI, and optical imaging. Scientific Reports. 2020; 10: 17324.

71. Goubran M, Leuze C, Hsueh B, Aswendt M, Ye L, Tian Q, et al. Multimodal image registration and connectivity analysis for integration of connectomic data from microscopy to MRI. Nature Communications. 2019; 10: 5504.

72. Cao Y, Vassantachart A, Ye JC, Yu C, Ruan D, Sheng K, et al. Automatic detection and segmentation of multiple brain metastases on magnetic resonance image using asymmetric UNet architecture. Phys Med Biol. 2021; 66: 015003.

73. Wang S, Pang X, de Keyzer F, Feng Y, Swinnen JV, Yu J, et al. AI-based MRI auto-segmentation of brain tumor in rodents, a multicenter study. Acta Neuropathologica Communications. 2023; 11: 11.

74. Karreman MA, Bauer AT, Solecki G, Berghoff AS, Mayer CD, Frey K, et al. Active Remodeling of Capillary Endothelium via Cancer Cell-Derived MMP9 Promotes Metastatic Brain Colonization. Cancer Res. 2023; 83: 1299–314.

75. Kovalchuk B, Berghoff AS, Karreman MA, Frey K, Piechutta M, Fischer M, et al. Nintedanib and a bi-specific anti-VEGF/Ang2 nanobody selectively prevent brain metastases of lung adenocarcinoma cells. Clin Exp Metastasis. 2020; 37: 637–48.

76. Tehranian C, Fankhauser L, Harter PN, Ratcliffe CDH, Zeiner PS, Messmer JM, et al. The PI3K/Akt/mTOR pathway as a preventive target in melanoma brain metastasis. Neuro Oncol. 2022; 24: 213–25.

77. Venkataramani V, Karreman MA, Nguyen LC, Tehranian C, Hebach N, Mayer CD, et al. Direct excitatory synapses between neurons and tumor cells drive brain metastatic seeding of breast cancer and melanoma. bioRxiv. 2024.

78. Capper D, Jones DTW, Sill M, Hovestadt V, Schrimpf D, Sturm D, et al. DNA methylation-based classification of central nervous system tumours. Nature. 2018; 555: 469–74.

79. Dondzillo A, Satzler K, Horstmann H, Altrock WD, Gundelfinger ED, Kuner T. Targeted three-dimensional immunohistochemistry reveals localization of presynaptic proteins Bassoon and Piccolo in the rat calyx of Held before and after the onset of hearing. J Comp Neurol. 2010; 518: 1008–29.

80. Kienast Y, von Baumgarten L, Fuhrmann M, Klinkert WE, Goldbrunner R, Herms J, et al. Real-time imaging reveals the single steps of brain metastasis formation. Nat Med. 2010; 16: 116–22.

81. Zitová B, Flusser J. Image registration methods: a survey. Image and Vision Computing. 2003; 21: 977–1000.

82. Rueckert D, Sonoda LI, Hayes C, Hill DL, Leach MO, Hawkes DJ. Nonrigid registration using free-form deformations: application to breast MR images. IEEE Trans Med Imaging. 1999; 18: 712–21.

83. Bookstein FL. Principal warps: thin-plate splines and the decomposition of deformations. IEEE Transactions on Pattern Analysis and Machine Intelligence. 1989; 11: 567–85.

84. Shinohara RT, Sweeney EM, Goldsmith J, Shiee N, Mateen FJ, Calabresi PA, et al. Statistical normalization techniques for magnetic resonance imaging. Neuroimage Clin. 2014; 6: 9–19.

85. Fernandez R, Moisy C. FijiRelax: Fast and noise-corrected estimation of MRI relaxation maps in 3D + t. Journal of Open Source Software. 2023; 8.

86. Yates SC, Groeneboom NE, Coello C, Lichtenthaler SF, Kuhn PH, Demuth HU, et al. QUINT: Workflow for Quantification and Spatial Analysis of Features in Histological Images From Rodent Brain. Front Neuroinform. 2019; 13: 75.

