## Supplementary Figures for "Voxel-accurate MRI-microscopy correlation enables AI-powered prediction of brain disease states"

### Supplementary material

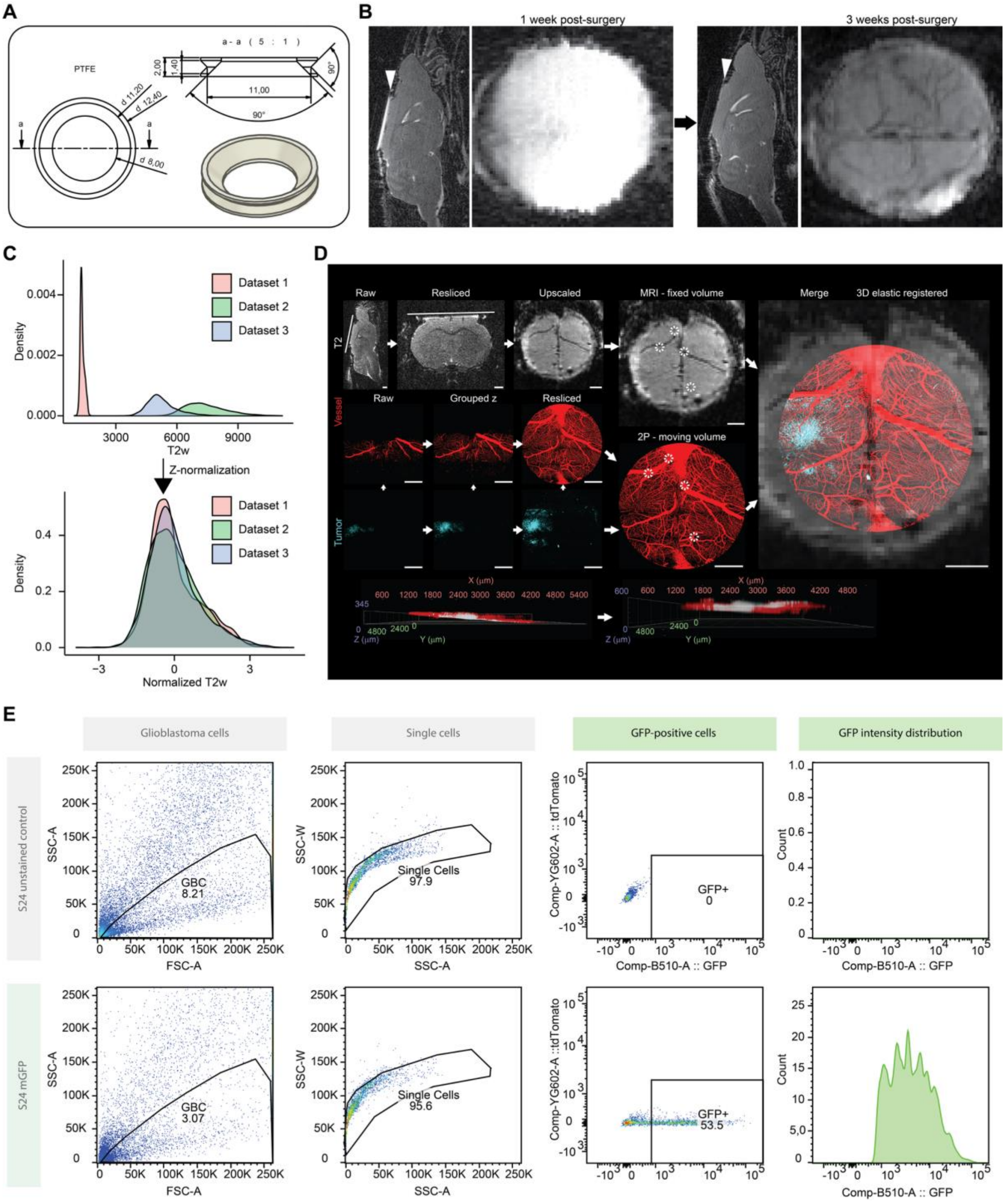


#### Figure S1 – *In vivo* BRIDGE workflow

**A,** Model of custom head fixation ring for 2P made out of Teflon (PTFE). Values in mm. **B,** *In vivo* MRI (T2w) of the same mouse 1 week and 3 weeks post-surgery. Arrowheads indicate cranial window glass. Scale bars: 1 mm. **C,** Density plots of MR voxel value distribution of different mice before (top) and after (bottom) z-normalization on healthy brain tissue MR voxel values (n = 3 mice, 3 datasets, 15447 voxels). **D,** Multistep workflow including reslicing MRI and 2P images, rescaling, and elastic 3D landmark registration in 3D Slicer using blood vessel landmarks - visualized in 2D. Scale bars: 1 mm. **E,** Gating strategies exemplary for sorting S24 mGFP (green, n = 100000 cells) with FACS using unstained controls (gray, n = 51359 cells).

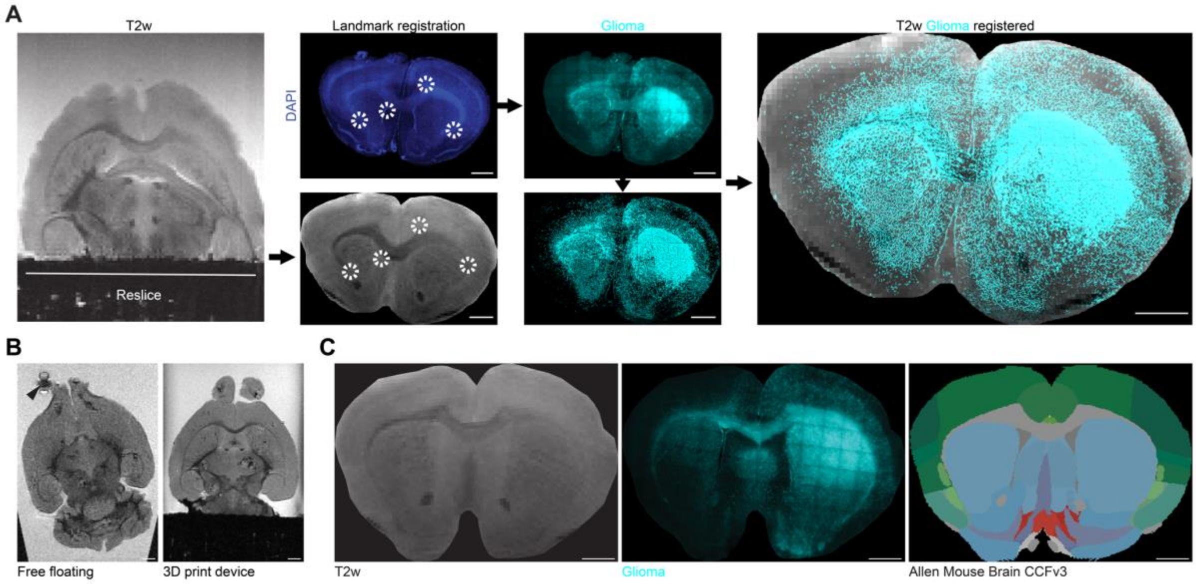


**Figure S2 – Bespoke 3D-printed device for enhanced *ex vivo* brain co-registration**

**A**, Computational integration of reslicing, MRI upscaling, and 3D landmark registration in an S24 mGFP PDX slice. DAPI (blue) and Nestin (cyan). Scale bars: 1 mm. **B,** Comparative *Ex vivo* T2*w imaging of Jimt1 xenograft mice brains acquired without (left) and with (right) 3D print device. Arrowhead depicts a gas bubble artifact. Scale bars: 1 mm. **C,** Slice alignment with Allen Mouse Brain CCFv3. Scale bars: 1 mm.

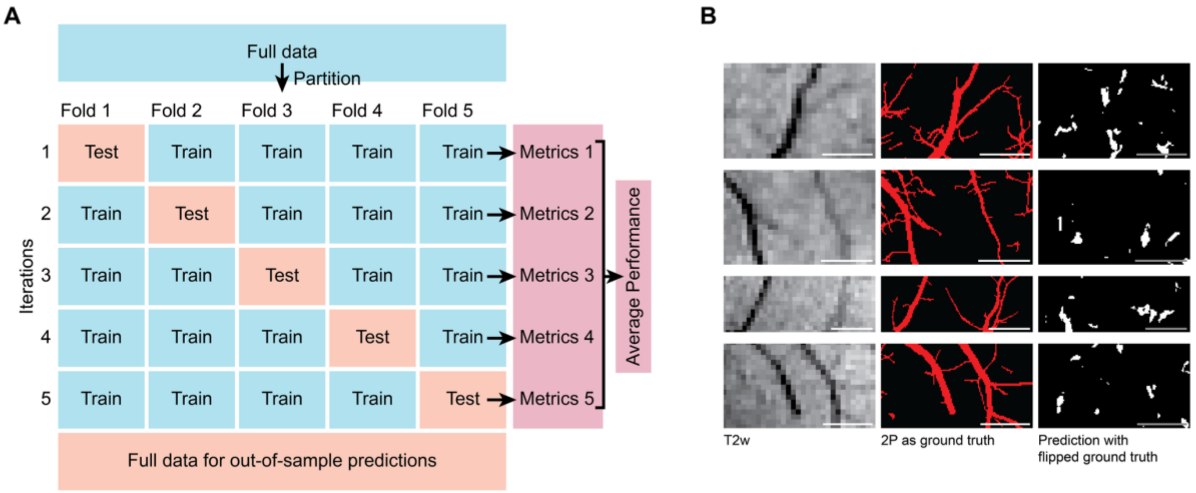


**Figure S3 – BRIDGE simplifies automatic segmentation of MR images using microscopy as ground truth**

**A,** Illustration of 5-fold cross-validation. **B,** Left: T2w input crops, middle: preprocessed 2P data as ground truth, right: predictions of training with vertically and horizontally flipped 2P data as ground truth to confuse the convolutional neural network. Scale bars: 1 mm.

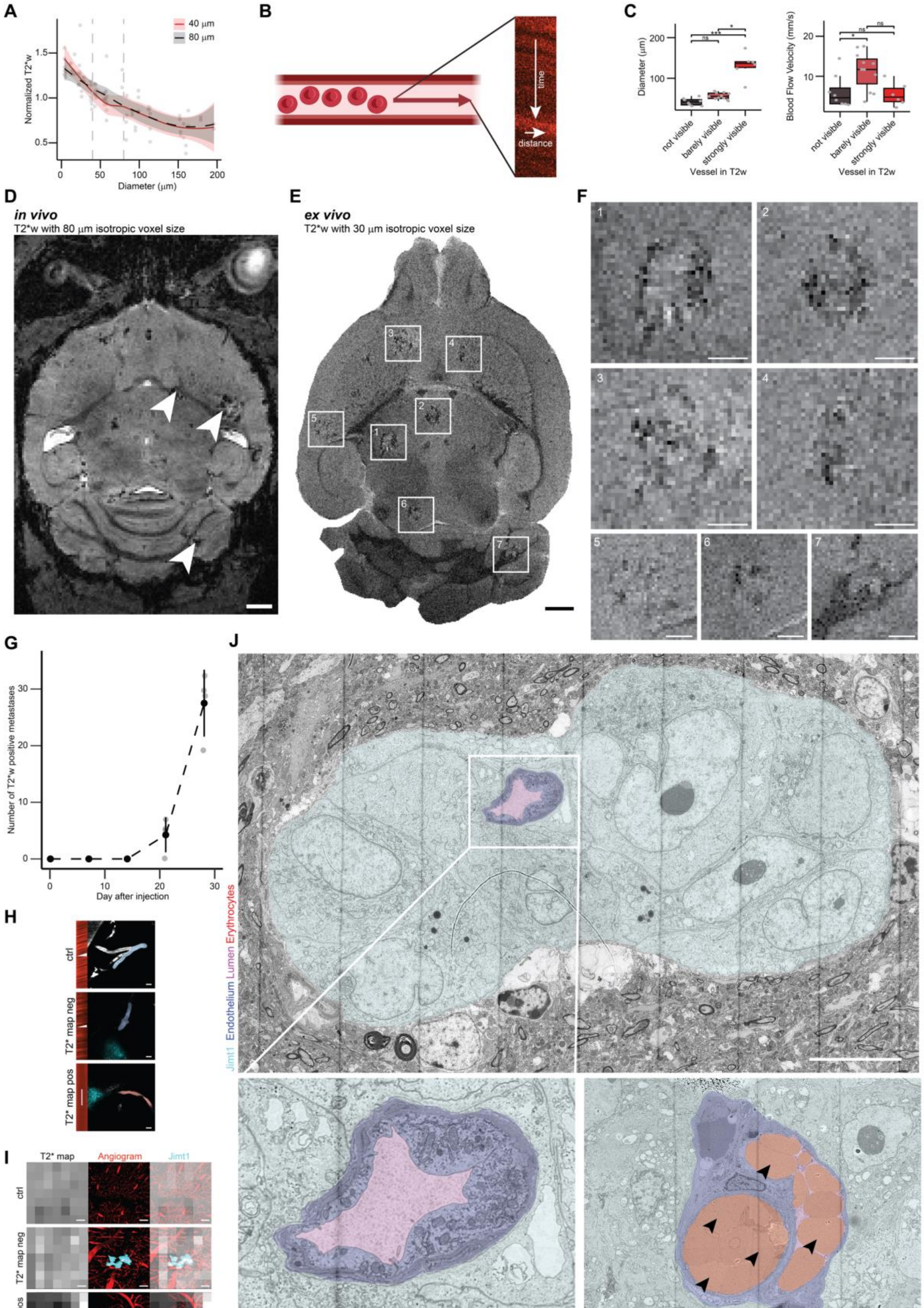
A
B
C
D
E
F
G
H
I
J
40 μm
80 μm
Normalized T2*w
Diameter (μm)
time
distance
Diameter (μm)
Blood Flow Velocity (mm/s)
not visible
barely visible
strongly visible
Vessel in T2w
ns
in vivo
T2*w with 80 μm isotropic voxel size
ex vivo
T2*w with 30 μm isotropic voxel size
Number of T2*w positive metastases
Day after injection
ctrl
T2* map neg
T2* map pos
T2* map
Angiogram
Jimt1
Jimt1 Endothelium Lumen Erythrocytes
No erythrostasis
Erythrostasis

**Figure S4 – Breast-cancer metastases show distinct T2*w positive lesions**

**A,** Line plot with smoothing function showing comparison of vessel diameters against T2*w intensities between MRI measurements at 40 µm and 80 µm resolution. Dashed lines indicate 40 µm (left) and 80 µm (right) (n = 40 vessel pairs in 1 mouse). **B,** Schematic of intravital blood flow measurement. **C,** Left: Diameter of vessels depending on visibility in T2w. Right: Blood flow velocity in vessels depending on visibility in T2w. Median with interquartile range (n = 2 mice; for T2w: not visible, n = 8 vessels; barely visible, n = 11 vessels; visible, n = 2 vessels; strongly visible, n = 4 vessels; Kruskal-Wallis test followed by Dunn-Bonferroni post-hoc test). **D,** *In vivo* T2*w sequence of Jimt1 breast cancer brain metastasis with isotropic voxel size of 80 µm. Arrowheads show metastases with T2*w hypointensities. Scale bar: 1 mm. **E,** *Ex vivo* high resolution T2*w sequence of Jimt1 breast cancer brain metastasis with isotropic voxel size of 30 µm. Squares show metastases with T2*w hypointensities. Scale bar: 1 mm. **F,** Larger illustrations of the outlined metastases from c. Scale bars: 300 µm. **G,** Temporal quantification of *in vivo* T2*w hypointense lesions in Jimt1 brains over 28 days (n = 4 mice, 20 datasets, 5 datasets per time point). **H,** Examples of blood flow velocity in control, T2*-negative lesion, and T2*-positive lesions. Scale bars: 10 µm. **I,** Examples of control, T2*-negative lesion, and T2*-positive lesions. Scale bars: 100 µm. **J,** Electron microscopy (EM) of perimetastatic capillaries showing a vessel with an empty lumen (top) and a vessel with erythrostasis (bottom). Arrowheads indicate erythrocytes. Jimt1 (cyan) and perimetastatic capillaries with endothelium (purple), erythrocytes (red) and lumen (pink). Scale bars: 10 µm.

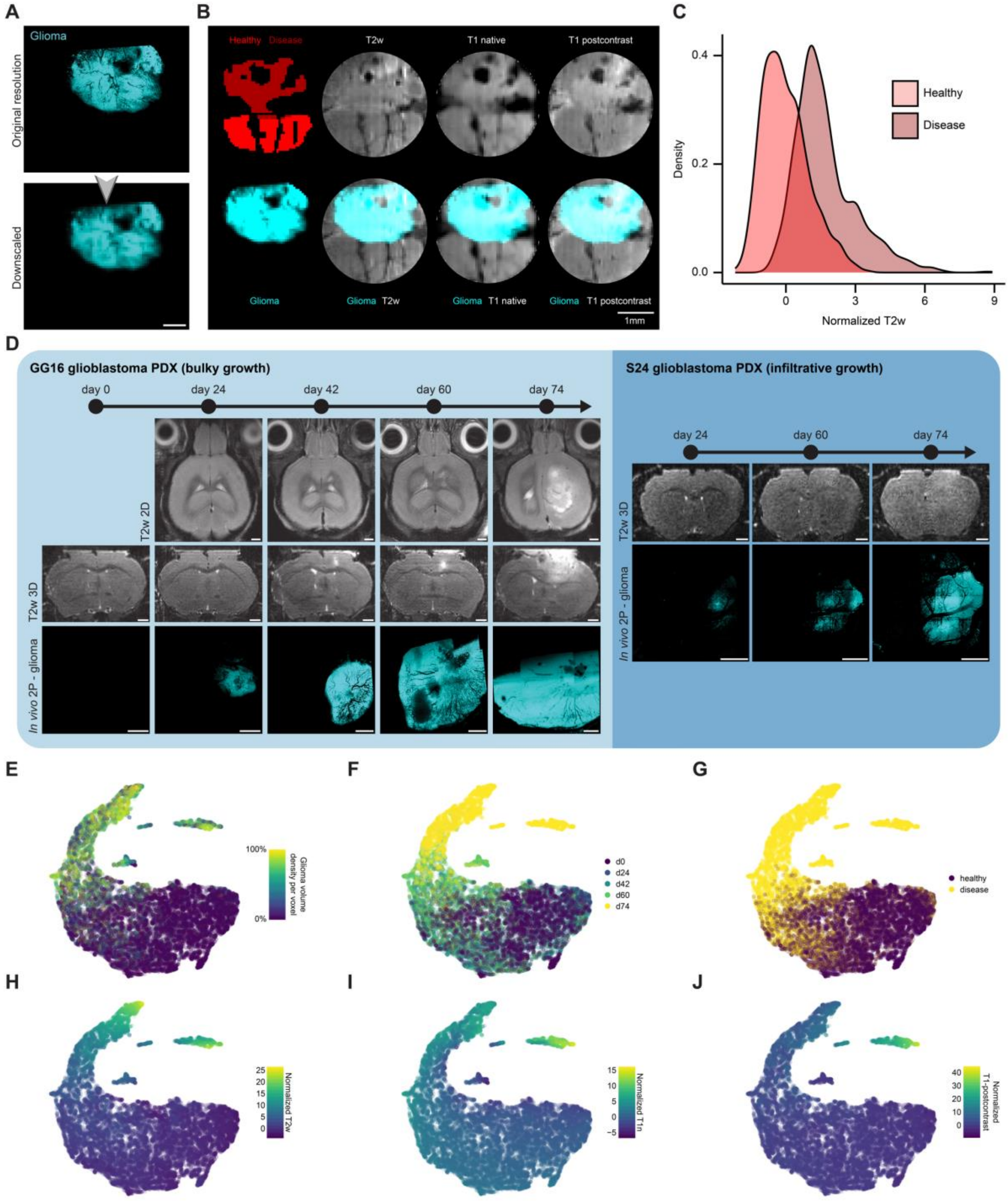


**Figure S5 – Voxel segmentation for correlative signal intensity measurements in glioblastoma**

**A,** Exemplary 2P image of GG16 with 2P resolution (1.18 µm) and downscaling to MR resolution (100 µm). Scale bars: 1 mm. **B,** 2P-MRI correlation of GG16 tumor (cyan) across multisequence MRI including T2, T1-native, and T1-postcontrast. Healthy tissue (red) and tumor (dark red). Scale bars: 1 mm. **C,** Density plot of normalized MR voxel grouped in “healthy” (red) and “disease” (dark red) of an exemplary dataset (day 42 after GG16 injection) (n = 1 dataset, 4007 voxels). **D,** Illustration of tumor growth in GG16 and S24 in MRI and 2P microscopy. Scale bars: 1 mm. **E-J,** UMAP dimension reduction of normalized T2w, T1-native, and T1-postcontrast data for GG16, labeled by glioma volume density (E), days after injection (F), mask categories (G), normalized T2w (H), normalized T1n (I) and normalized T1-postcontrast (J) (n = 4 mice, 13 datasets, 27555 voxels).

**Movie S1 – Longitudinality, multimodality and precision of the BRIDGE co-registration pipeline *in vivo***

Visualization of the different sequences on which 2P microscopy is registered based on unique vessel branches. Even small vessels show correspondence between MRI and 2P microscopy. Time course and tumor growth follow.

**Movie S2 – Bridging scales from MRI to super-resolution microscopy *ex vivo***

The 3D representation of a brain mask segmented in MRI images is followed by a slice of the *ex vivo* T2w sequence. Microscopy was registered on this slice. Expansion microscopy was performed to see individual NGS.

**Movie S3 – Clinical translation bridges scales from MRI to super-resolution microscopy in human glioblastoma tissue**

Neuronavigation mask is followed by the *ex vivo* T2w sequence. Microscopy was co-registered. In the following, expansion microscopy was performed to visualize glioma cells in 3D.

**Table S1 – Tumor cell line classification**

**Table S2 – *In vivo* MRI sequences**

**Table S3 – *Ex vivo* MRI sequences**

**Table S4 – Reagents**

**Table S5 – Organisms and strains**

**Table S6 – Devices**

**Table S7 – Programs**
